# Cross-platform genetic discovery of small molecule products of metabolism and application to clinical outcomes

**DOI:** 10.1101/2020.02.03.932541

**Authors:** Luca A. Lotta, Maik Pietzner, Isobel D. Stewart, Laura B.L. Wittemans, Chen Li, Roberto Bonelli, Johannes Raffler, Emma K. Biggs, Clare Oliver-Williams, Victoria P.W. Auyeung, Jian’an Luan, Eleanor Wheeler, Ellie Paige, Praveen Surendran, Gregory A. Michelotti, Robert A. Scott, Stephen Burgess, Verena Zuber, Eleanor Sanderson, Albert Koulman, Fumiaki Imamura, Nita G. Forouhi, Kay-Tee Khaw, MacTel Consortium, Julian L. Griffin, Angela M. Wood, Gabi Kastenmüller, John Danesh, Adam S. Butterworth, Fiona M. Gribble, Frank Reimann, Melanie Bahlo, Eric Fauman, Nicholas J. Wareham, Claudia Langenberg

**Affiliations:** MRC Epidemiology Unit, University of Cambridge, Cambridge, UK; The Big Data Institute, Li Ka Shing Centre for Health Information and Discovery, University of Oxford; Population Health and Immunity Division, The Walter and Eliza Hall Institute of Medical Research, Parkville, Australia; Department of Medical Biology, The University of Melbourne, Parkville, Australia; Institute of Computational Biology, Helmholtz Zentrum München – German Research Center for Environmental Health, Neuherberg, Germany; Metabolic Research Laboratories, University of Cambridge, Cambridge, United Kingdom; British Heart Foundation Cardiovascular Epidemiology Unit, Department of Public Health and Primary Care, University of Cambridge, Cambridge, UK; Homerton College, University of Cambridge, Cambridge, UK; National Centre for Epidemiology and Population Health, The Australian National University, Canberra, Australia; British Heart Foundation Centre of Research Excellence, University of Cambridge, Cambridge, UK; Health Data Research UK Cambridge, Wellcome Genome Campus and University of Cambridge, Cambridge, UK; Rutherford Fund Fellow, Department of Public Health and Primary Care, University of Cambridge, UK; Metabolon Inc, Durham, North Carolina USA; MRC Biostatistics Unit, University of Cambridge, Cambridge, United Kingdom; Department of Public Health and Primary Care, University of Cambridge, Cambridge, United Kingdom; Department of Epidemiology and Biostatistics, Imperial College London, UK; MRC Integrative Epidemiology Unit, Bristol Medical School, University of Bristol, UK; NIHR BRC Nutritional Biomarker Laboratory, University of Cambridge, UK; Biomolecular Medicine, Department of Metabolism, Digestion and Reproduction, Imperial College London, UK; National Institute for Health Research Blood and Transplant Research Unit in Donor Health and Genomics, University of Cambridge, Cambridge, UK; The Alan Turing Institute, London, UK; National Institute for Health Research Cambridge Biomedical Research Centre, University of Cambridge and Cambridge University Hospitals, Cambridge, UK; Department of Human Genetics, Wellcome Sanger Institute, Hinxton, UK; Internal Medicine Research Unit, Pfizer Worldwide Research, Cambridge, MA 02142, USA

## Abstract

Circulating levels of small molecules or metabolites are highly heritable, but the impact of genetic differences in metabolism on human health is not well understood. In this cross-platform, genome-wide meta-analysis of 174 metabolite levels across six cohorts including up to 86,507 participants (70% unpublished data), we identify 499 (362 novel) genome-wide significant associations (p<4.9×10^-10^) at 144 (94 novel) genomic regions. We show that inheritance of blood metabolite levels in the general population is characterized by pleiotropy, allelic heterogeneity, rare and common variants with large effects, non-linear associations, and enrichment for nonsynonymous variation in transporter and enzyme encoding genes. The majority of identified genes are known to be involved in biochemical processes regulating metabolite levels and to cause monogenic inborn errors of metabolism linked to specific metabolites, such as *ASNS* (rs17345286, MAF=0.27) and asparagine levels. We illustrate the influence of metabolite-associated variants on human health including a shared signal at GLP2R (p.Asp470Asn) associated with higher citrulline levels, body mass index, fasting glucose-dependent insulinotropic peptide and type 2 diabetes risk, and demonstrate beta-arrestin signalling as the underlying mechanism in cellular models. We link genetically-higher serine levels to a 95% reduction in the likelihood of developing macular telangiectasia type 2 [odds ratio (95% confidence interval) per standard deviation higher levels 0.05 (0.03-0.08; p=9.5×10^-30^)]. We further demonstrate the predictive value of genetic variants identified for serine or glycine levels for this rare and difficult to diagnose degenerative retinal disease [area under the receiver operating characteristic curve: 0.73 (95% confidence interval: 0.70-0.75)], for which low serine availability, through generation of deoxysphingolipids, has recently been shown to be causally relevant. These results show that integration of human genomic variation with circulating small molecule data obtained across different measurement platforms enables efficient discovery of genetic regulators of human metabolism and translation into clinical insights.

## Introduction

Metabolites are small molecules that reflect biological processes and are widely measured in clinical medicine as diagnostic, prognostic or treatment response biomarkers^1^. Blood levels of metabolites are highly heritable with twin studies reporting a median explained variance in plasma levels of 6.9% and maximum of 50% depending on the metabolite^2,3^. Several earlier studies have started to characterise the genetic architecture of metabolite variation in the general population^2–10^, but been limited in size and scope by focussing on metabolites assessed using a single method. Integration of genetic association results for metabolites measured on different platforms can help maximise the power for a given metabolite and provide a more refined understanding of genetic influences on blood metabolite levels and human physiology.

To identify genomic regions regulating metabolite levels and systematically study their relevance for disease, we designed and conducted a cross-platform meta-analysis of genetic effects on levels of 174 blood metabolites measured in large-scale population-based studies. We included metabolites covered by the targeted Biocrates AbsoluteIDQ^™^ p180 platform and measured in the Fenland Study. We integrated unpublished data for any of these metabolites that were covered by the Nightingale (^1^H-NMR, Interval Study) or Metabolon (Discovery HD4™, EPIC-Norfolk and Interval Studies) platforms, or had previously been reported^2,4,5^. The focus on this targeted set of ‘platform-specific’ metabolites enabled us to clearly map metabolites across platforms and maximise the sample size for each of the 174 metabolites for this proof of concept cross-platform GWAS study. To facilitate rapid sharing of our results, we developed a webserver (https://omicscience.org/apps/crossplatform/) that allows flexible interrogation of our results.

## Results

### Associations with blood metabolites at 144 genomic regions

Genome-wide meta-analyses were conducted for 174 metabolites from 7 biochemical classes (i.e. amino acids, biogenic amines, acylcarnitines, lyso-phosphatidylcholines, phosphatidylcholines, sphingomyelins and hexose) commonly measured using the Biocrates p180 kit in up to 86,507 individuals, contributing over 3.7 million individual-metabolite data points (70% from unpublished studies; **Fig. 1**). For each of the 174 metabolites, this was the largest genome-wide association analyses (GWAS) to date, with at least a doubling of sample size (**Fig. 1A**). Sample sizes ranged from 8,569 to 86,507 individuals for metabolites depending on the platform used in each contributing study. Using GWAS analyses we estimated the association of up to 10.2 million single nucleotide variants with a minor allele frequency (MAF) >0.5%, including 6.1 million with MAF ≥ 5%.

**Figure 1.**
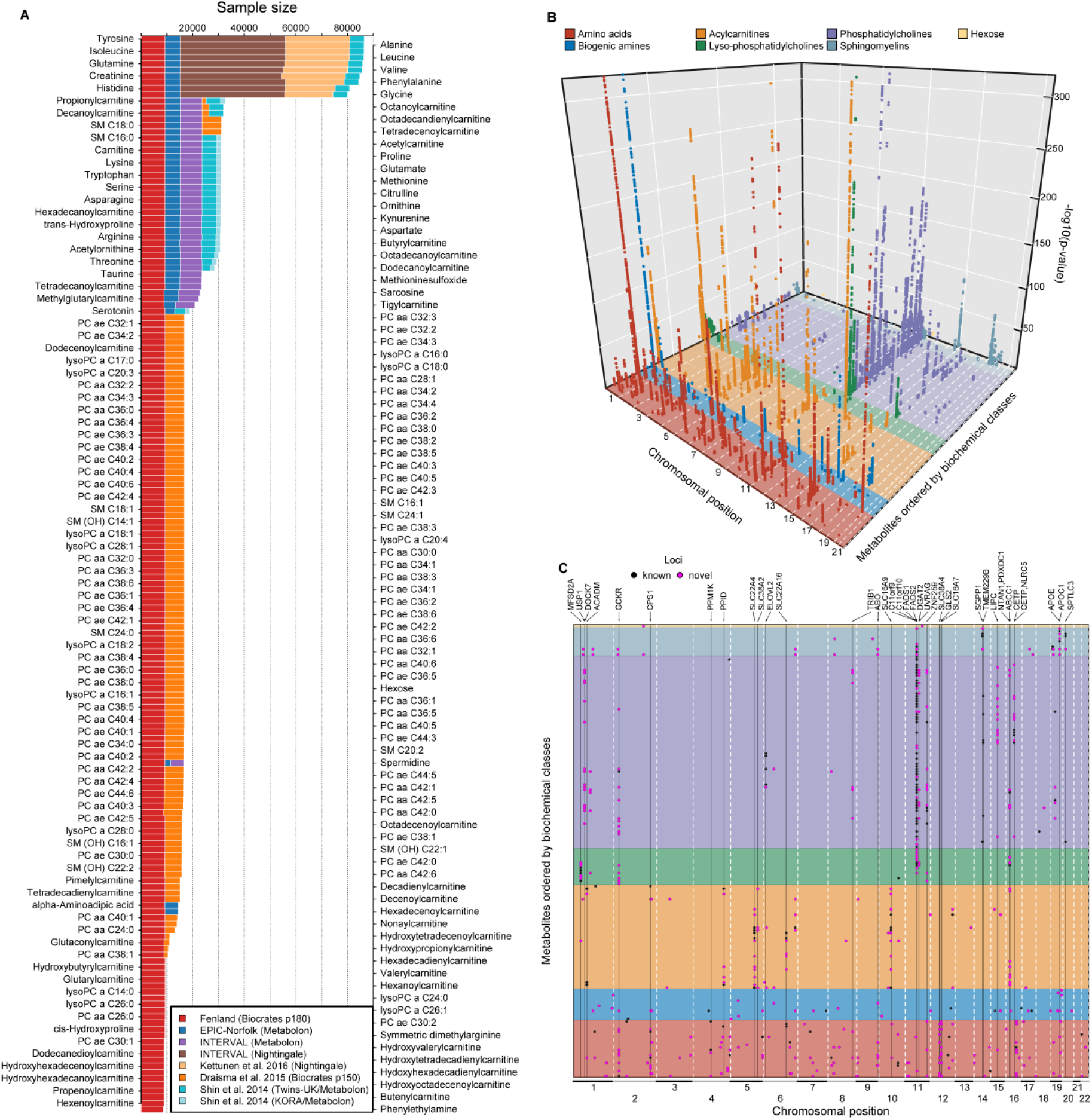
**A** Sample size by contributing study and technique for each of the 174 metabolites included. **B** A three-dimensional Manhattan plot displaying chromosomal position (x-axis) of significant associations (p <4.9×10^-10^, z-axis) across all metabolites (y-axis). Colours indicate metabolite groups. **C** A top view of the 3D-Manhattan plot. Dots indicate significantly associated loci. Colours indicate novelty of metabolite - locus associations. Loci with indication for pleiotropy have been annotated.

We identified 499 variant-metabolite associations (362 novel) from 144 loci (94 novel) at a metabolome-adjusted genome-wide significance threshold of p<4.9×10^-10^ (correcting the usual GWAS-threshold, p<5×10^-8^, for 102 principal components explaining 95% of the variance in metabolite levels using principal component analysis; **Fig. 1**). The vast majority of these associations were consistent across studies and measurement platforms [median I^2^: 26.8 (interquartile range: 0 – 70.1) for 465 associations with at least two contributing studies] (**Supplementary Tab. S1-2**). To identify possible sources of heterogeneity, we investigated the influence of differences by cohort, measurement platform, metabolite class, and association strength in a joint meta-regression model (**Supplementary Tab. S3**). This showed that heterogeneity was mainly due to the overall strength of the signal, i.e. associations with higher z-scores showed greater heterogeneity (p<1.05×10^-9^). However, the majority of these statistically heterogeneous associations were directionally consistent and nominally significant across and within each stratum for 146 of 170 associations with a z-score > 10, demonstrating the feasibility of pooling association estimates across metabolomics platforms for the purpose of genetic discovery. Genetic variants at the *NLRP12* locus, e.g. rs4632248, were a notable exception with large estimates of heterogeneity (I^2^>90%). The *NLRP12* locus is known to affect the monocyte count^11^ and has been shown to have pleiotropic effects on the plasma proteome in the INTERVAL study^12^. Monocytes, or at least a subpopulation subsumed under this cell count measure, release a wide variety of biomolecules upon activation or may die during the sample handling process and hence releasing intracellular biomolecules, such as taurine^13^, into the plasma. In brief, one specific source of heterogeneity in mGWAS associations might relate to sample handling differences across studies.

This highlights the utility of our genetic cross-platform approach to maximise power for a given metabolite, substantially extending previous efforts for any given metabolite^14^. Previously reported associations from platform-specific studies were also found to generally be consistent in our cross-platform meta-analysis (**Supplementary Tab. S2;** https://omicscience.org/apps/crossplatform/).

### Insights in the genetic architecture of metabolite levels

We identified a median of 2 (range: 1-67, **Fig. 2A**) associated metabolites for each locus and a median of 3 (range: 1-20, **Fig. 2B**) locus associations for each metabolite, reflecting pleiotropy and the extensive contribution of genetic loci to circulating metabolite levels. The number of associations was proportional to the estimated heritability and the sample size of the meta-analysis for a given trait (**Fig. 2C**).

**Figure 2.**
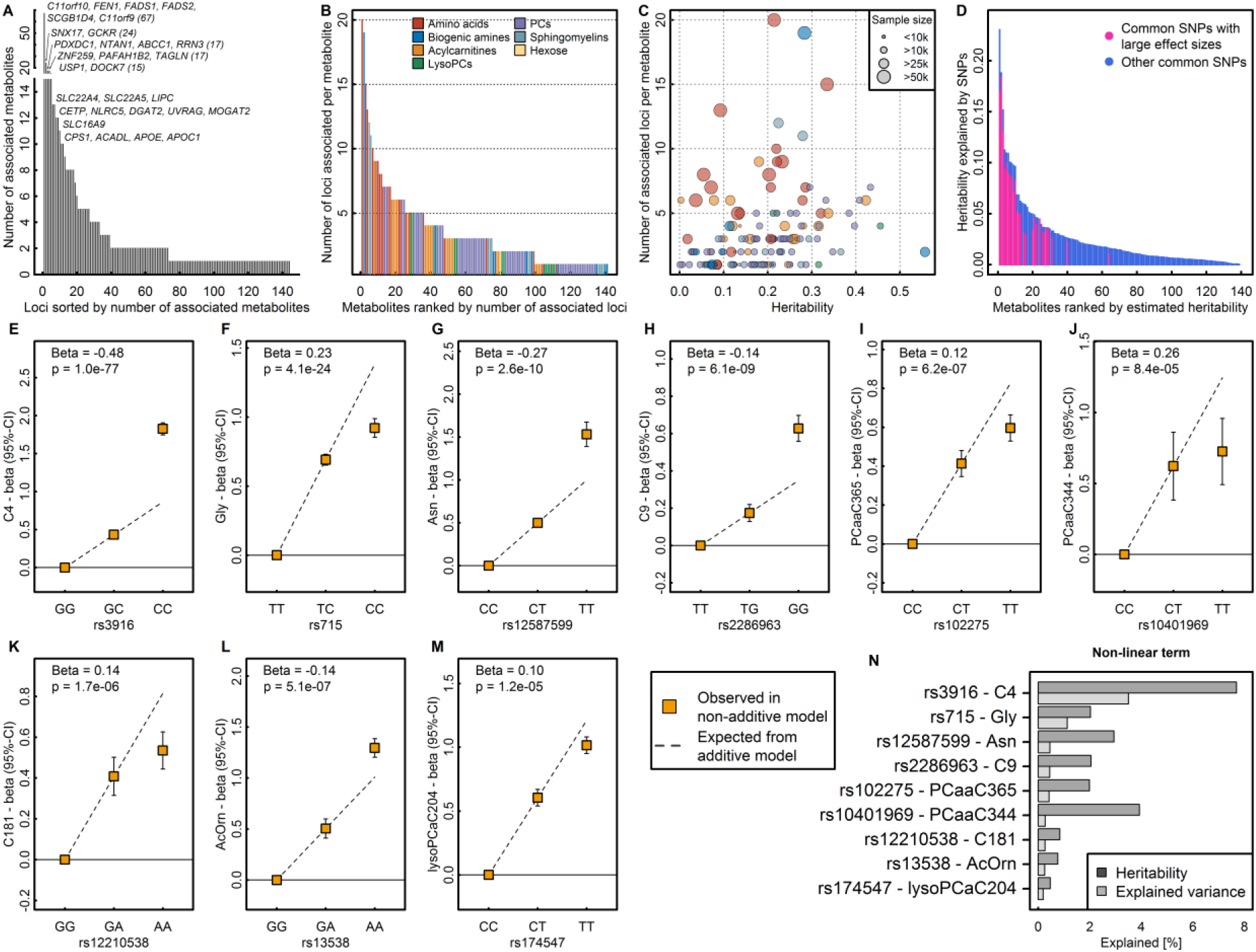
**A** Distribution of pleiotropy, i.e. number of associated metabolites, among loci identified in the present study. **B** Distribution of polygenicity of metabolites, i.e. number of identified loci for each metabolite under investigation. **C** Scatterplot comparing the estimated heritability of each metabolite against the number of associated loci. Size of the dots indicates samples sizes. **D** Heritability estimates for single metabolites. Colours indicate the proportion of heritability attributed to single nucleotide polymorphisms (SNPs) with large effect sizes (β>0.25 per allele). **E – M** SNP – metabolite association with indication of non-additive effects. Beta is an estimate from the departure of linearity. **N** Barplot showing the increase in heritability and explained variance for each SNP – metabolite pair when including non-additive effects.

We applied a multi-trait statistical colocalisation method^15^ and identified between 1-30 (median: 2) metabolites that did not meet the discovery p-value threshold, but showed high posterior probability (>75%) of a shared genetic signal for 49 out of the 144 loci (**Supplemental Fig. S1**). Two distinct variants (rs2414577 and rs261334) nearby *LIPC* showed the largest gain in additionally associated metabolites, in line with previous reports of extensive pleiotropy and allelic heterogeneity at this locus^9^. We note that a low posterior probability for the alignment of multiple metabolites at other loci might be explained by the presence of multiple causal variants shared across multiple metabolites.

To systematically classify pleiotropic variants taking into account the correlation structure among metabolites we derived a data-driven metabolic network and performed community detection (see **Methods** and **Supplemental Fig. S2**). A total of 129 (60.5%) of 214 variants (associated with at least two metabolites at p<5×10^-8^) were associated with metabolites from at least two of the 14 communities (range: 2 – 11; **Supplemental Fig. S2**), i.e. showed evidence for ‘horizontal’ or broad pleiotropy. The most extreme variants included those near *FADS1* (e.g. rs17455) associated with 61 metabolites across 11 communities at p<5×10^-8^. In contrast, rs2638315 (likely tagging a missense variant rs2657879 at *GLS2*) was associated with nine metabolites within a single community and would therefore be considered as ‘horizontal pleiotropic’ for a well-defined group of correlated metabolites (**Supplemental Fig. S2**).

Similar to what is routinely observed in GWAS literature, effect size estimates increased with decreasing minor allele frequency (MAF) (**Fig. 3A**). However, there were 26 associations (**Tab. 1**) for common lead variants with per-allele differences in metabolites levels greater than 0.25 standard deviations (SD), a per-allele effect size that is >3-fold larger than the strongest common variants associated with SDs of body mass index at the *FTO* locus.

**Figure 3.**
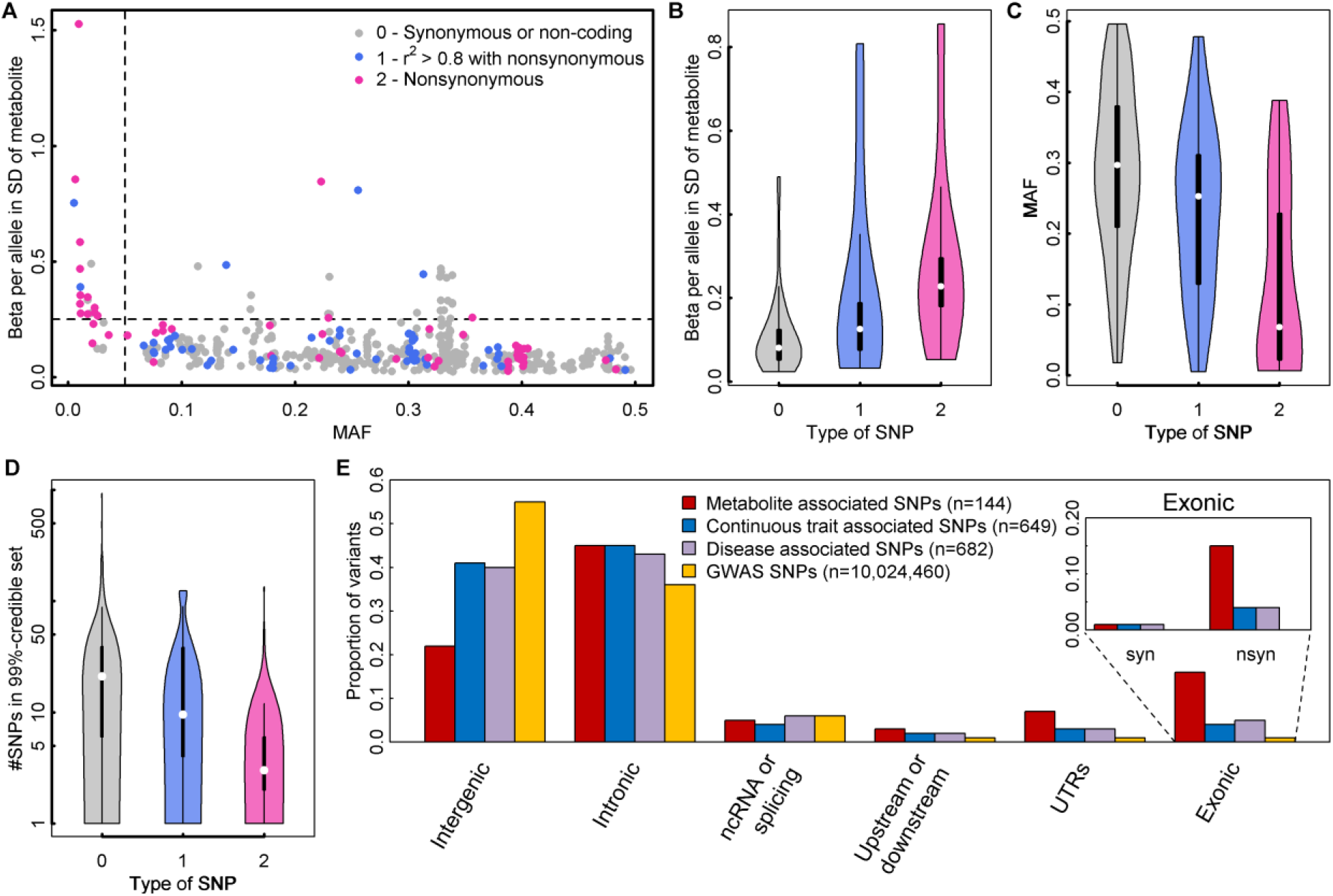
**A** Scatterplot comparing the minor allele frequencies (MAF) of associated variants with effect estimates from linear regression models (N loci=499). Colours indicate possible functional consequences of each variant: maroon – nonsynonymous variant; blue – in strong LD (r^2^>0.8) with a nonsynonymous variant and grey otherwise. **B-D** Distribution of effect sizes (B), allele frequencies (C), and width of credible sets (D) based on the type of single nucleotide polymorphism (SNP) (0 – non-coding or synonymous, 1 – in strong LD with nonsynonymous, 2 – nonsynonymous). **E** Distribution of functional annotations of metabolite associated variants (red), trait-associated variants (blue – continuous, purple – diseases) obtained from the GWAS catalogue, and all SNPs included in the present genome-wide association studies. The inlet for exonic variants distinguishes between synonymous (syn) and nonsynonymous variants (nsyn).

**Table 1.**
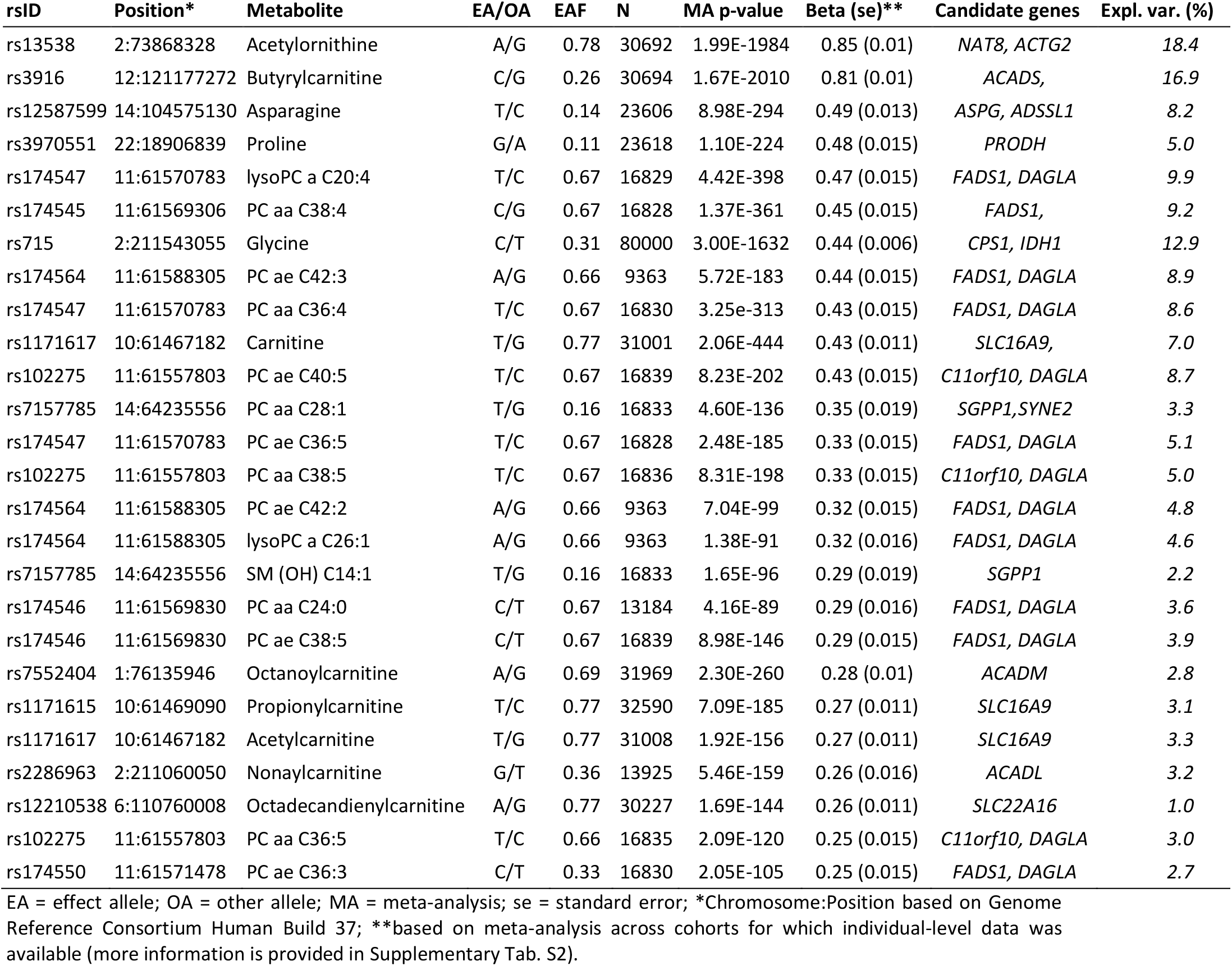
Genomic loci with effect sizes larger than 0.25 units in standard deviation of metabolite levels per allele.

Variants identified in this study explained up to 23% of the variance (median: 1.4%; interquartile range: 0.5% - 2.8%) and up to 99.8% of the chip-based heritability (median 9.2%; interquartile range: 4.7% - 17.1%) for the 141 metabolites with at least one genetic association (**Fig. 2D**). The 26 common variants with large effect sizes (>0.25 SD per allele) were identified for metabolites with higher heritability (**Fig. 2D**) and accounted for up to 74% of the heritability explained in those metabolites.

GWAS analyses generally assume a linear relationship between genotypes and phenotypes, i.e. an additive dose-response model. The identification of several metabolite-associated variants with large effect sizes and availability of individual-level data in the Fenland cohort allowed us to test whether the metabolite-associated variants showed evidence of deviation from a linear model. Of 499 associations tested, 9 showed evidence of departure from a linear association (**Fig. 2E-M**). Modelling actual genotypes rather than assuming ‘additive’ linear associations in these instances explained a median of 7.4% more (range: 1.4-15.2%) of the heritability in metabolite levels (**Fig. 2N**). Associations better described by an autosomal recessive or dominant model of inheritance might be the most likely explanation for this. Variant rs3916, for example, which showed a more than additive positive effect on butyrylcarnitine, is in perfect LD with a missense variant within *ACADS* (rs1799958, MAF=26%), which encodes for short-chain acyl-CoA dehydrogenase (SCAD). SCAD deficiency is an autosomal recessive disease diagnosed by elevated butyrylcarnitine concentrations in blood and homozygeous carrier status for established pathogenic variants^16^.

In 61 of the 499 associations the lead association signal was a nonsynonymous variant, a 40-fold enrichment compared to what would be expected by chance given the annotation of ascertained genetic variants (two-tailed binomial test, p=5×10^-30^, **Fig. 3D**). For a further 59 associations, the lead variant was in high LD with a nonsynonymous variant (r^2^>0.8). Lead variants that were nonsynonymous, or variants in high LD with a nonsynonymous variant, generally had lower MAF, larger effect sizes, and smaller 99%-credible sets (**Supplemental Tab. S4**) than variants that were not in these categories (**Fig 3B-D**).

We identified 22 loci harbouring two (n=21) or three (n=1) independent signals, i.e. different plasma metabolites were associated with distinct genetic variants within the same genomic region (**Supplementary Tab. S2**). For six regions, our two different annotations approaches assigned only one causal gene (see below and **Methods**), including *ACADM, GLDC, ARG1, MARCH8, SLC7A2*, and *LIPC* (**Supplementary Tab. S2**). We found evidence that allelic heterogeneity, i.e. conditionally independent variants at a locus for a specific metabolite, explains the association pattern at 3 of those loci (*ACADM, ARG1*, and *LIPC*; **Supplementary Tab. S5**). We identified another 16 loci harbouring at least one (range: 2–6) additional conditionally independent variant(s) in exact conditional analyses (see **Methods**, **Supplementary Tab. S5**).

### Effector genes, tissues, pathways

We used two complementary strategies to prioritize likely causal genes for the observed associations: (1) a hypothesis-free genetic approach based on physical distance, genomic annotation and integration of expression quantitative trait loci (eQTLs) to prioritize genes in a systematic and standardised way (see **Methods**), and (2) a biological knowledge-based approach integrating existing knowledge about specific metabolites or related pathways to identify biologically plausible candidate genes from the 20 genes closest to the lead variant (**Fig. 4A**). Using the hypothesis-free genetic approach, we identified 249 unique likely causal genes for the 499 associations, with at least one gene per association and some genes prioritized as likely causal for multiple metabolite associations. The knowledge-based approach identified 130 biologically plausible genes for 349 out of 499 associations. We asked whether the hypothesis-free genetic approach identified biologically plausible genes (prioritized by strategy 2) more often than expected by chance. Amongst 9,980 possible gene-metabolite pairs (20 genes x 499 associations), 420 (4.2%) were biologically plausible, condensed to 350 gene(s)-metabolite assignments after accounting for overlapping annotations. Of the latter, 126 pairs (36%) were identical to genetically-prioritized gene-metabolite pairs, representing a significant enrichment of biologically plausible genes among those prioritised by the hypothesis-free algorithm (~8-fold more than expected by chance; two-tailed binomial test, p=2.3×10^-80^; **Fig. 4B**). Among the consistently assigned genes between both approaches, assignment of the nearest gene (124 times out of 126, χ^2^-test, p<2.5×10^-45^) was the strongest shared factor, as might be expected, followed by being (or in LD with) a missense variant (R^2^>0.8, 30 times out of 126, χ^2^-test, p<1.3×10^-07^) and only a minor contribution of eQTL data (20 times out of 126, χ^2^-test, p<0.001). Over 70% of genetically prioritized genes were enzymes or transporters (**Fig. 4C**). Inconsistencies between the approaches might be explained by non-consideration of information on biological pathways in the hypothesis-free genetic approach, as well as variants acting more distal to the biological determinants of plasma metabolite levels not being considered in the knowledge-based approach. The missense variant rs1260326 within *GCKR*, for example, colocalised with 49 metabolites across diverse biochemical classes (**Supplemental Fig. S1**) and likely confers it effects on glucose metabolism through impaired inhibition of glucokinase by glucokinase regulatory protein and might hence be considered as putative causal candidate by the knowledge-driven approach for plasma glucose only. However, impairments in glucose metabolism result in numerous downstream consequences including more distal metabolic branches such as amino acid and lipid metabolism.

**Figure 4.**
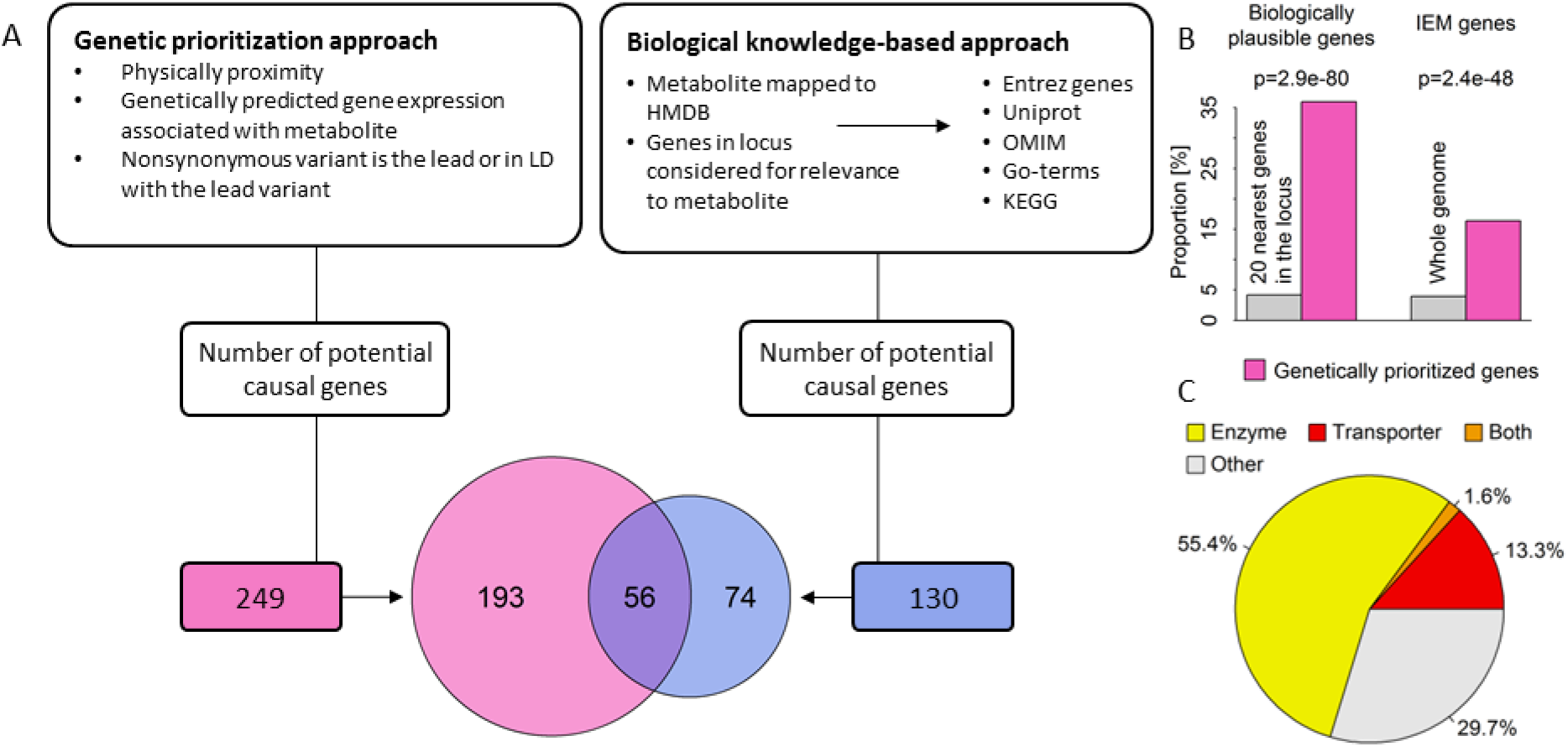
**A** Comparison between the hypothesis-free genetically prioritized versus biologically plausible approaches used in the present study to assign candidate genes to metabolite associated single nucleotide polymorphisms. The Venn-diagram displays the overlap between both approaches. **B** Enrichment of genetically prioritized genes among biologically plausible or genes linked to inborn errors of metabolism (IEM). **C** Proportion of genetically prioritized genes encoding for either enzymes or transporters.

In addition to being enriched in genes previously implicated in the biology of these metabolites, the genetically prioritized genes were also enriched in genes known for mutations to cause rare inborn errors of metabolism (IEMs), i.e. monogenic defects in the metabolism of small molecules with very specific metabolite changes (**Fig. 4B**).

Integrating GWAS statistics across cohorts and platforms allowed us to identify three genes that have never been associated with any metabolite level so far. At the *CERS6* locus, rs4143279 associates with levels of sphingomyelin (d18:1/16:0) (p = 4.2×10^-10^). *CERS6* encodes a ceramide synthase facilitating formation of ceramide, a precursor of sphingomyelins^17^. At the *ASNS* locus, rs17345286 associates with levels of asparagine (p = 4.7×10^-20^). The lead variant is in high LD (R^2^=1) with a missense mutation in *ASNS* (rs1049674, p.Val210Glu). *ASNS* encodes an asparagine synthase^18^. Finally, at the *SLC43A1* locus, rs2649667 associates with levels of phenylalanine (p = 3.6×10^-13^). *SLC43A1* encodes a liver-enriched transporter of large neutral amino acids, including phenylalanine^19^.

### Insights into the causes of common and rare diseases from metabolite-associated loci

The phenotypic consequences of metabolite-associated variants are currently not well characterized. Below, we investigate the contribution of individual loci and polygenic predisposition associated with differences in metabolite levels to the risk of common and rare diseases.

### A citrulline-raising functional variant in GLP2R increases type 2 diabetes risk

Because several of the metabolites captured in this GWAS have been associated with incident type 2 diabetes (T2D), we sought to investigate whether the association between metabolite-associated loci and diabetes could provide insights into underlying pathophysiologic mechanisms. Using estimates of effect for association with T2D based on a meta-analysis of 80,983 cases and 842,909 controls (see **Methods**), we observed a significant enrichment for associations with type 2 diabetes (p-value=2.8×10^-7^) of metabolite-associated variants compared to a matched control set of variants (**Fig. 5A**).

**Figure 5.**
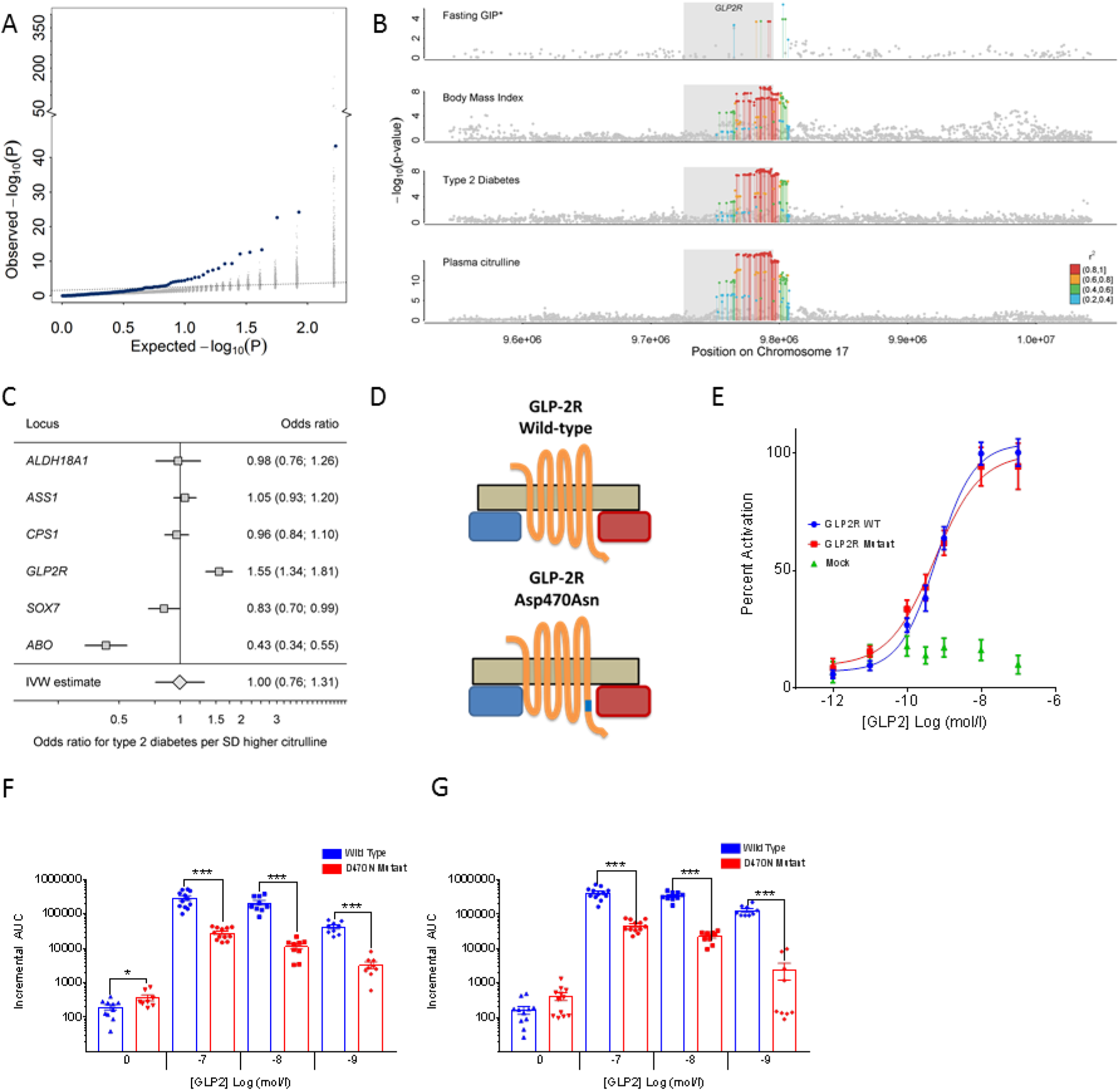
**A** Enrichment of associations with type 2 diabetes (T2D: 80,983 cases, 842,909 controls) among metabolite-associated SNPs. Blue dots indicate metabolite-SNPs and grey dots indicate a random selection of matched control SNPs. **B** Regional association plots for plasma citrulline, type 2 diabetes, body mass index, and fasting levels of glucose-dependent insulinotropic peptide (GIP) focussing on the *GLPR2* gene. Variants are coloured based on linkage disequilibrium with the lead variant (rs17681684) for plasma citrulline. *Summary statistics for GIP were obtained from the more densely genotyped study included in Almgren et al.^24^ (to increase coverage of genetic variants for multi-trait colocalisation). **C** Individual association summary statistics for all citrulline associated SNPs (coded by the citrulline increasing allele) for T2D and an inverse-variance weighted (IVW) estimate pooling all effects. **D** Schematic sketch for the location of the missense variant induces amino acid substitution in the glucagon-like peptide-2 receptor (GLP2R). **E** GLP-2 dose response curves in cAMP assay for GLP2R wild-type and mutant receptors. The dose response curves of cAMP stimulation by GLP-2 in CHO K1 cells transiently transfected with either GLP2R wild-type or mutant constructs. Data were normalised to the wild-type maximal and minimal response, with 100% being GLP-2 maximal stimulation of the wild-type GLP2R, and 0% being wild-type GLP2R cells with buffer only. Mean ± standard errors are presented (n=4).**F-G** Summary of wild-type and mutant GLP2R beta-arrestin 1 and beta-arrestin 2 responses. Area under the curve (AUC) summary data (n=3-4) displayed for beta-arrestin 1 recruitment (E) and beta-arrestin 2 recruitment (F). AUCs were calculated using the 5 minutes prior to ligand addition as the baseline value. Mean ± standard errors are presented. Normal distribution of log10 transformed data was determined by the D’Agostino & Pearson normality test. Following this statistical significance was assessed by one-way ANOVA with post hoc Bonferroni test. ***p<0.001, *p<0.05.

Amongst the diabetes- and metabolite-associated loci was a missense p.Asp470Asn (rs17681684) variant in the *GLP2R* gene encoding the receptor for glucagon-like peptide 2, a 33 amino acid peptide hormone encoded by the proglucagon gene (*GCG*) that stimulates the growth of intestinal tissue. Common variants at *GLP2R* are associated with an increased risk of T2D^20^. The previously reported lead variant for T2D (rs78761021) is in high LD (r^2^>0.87) with our lead citrulline association signal at *GLP2R* (rs17681684), which was associated with a 4% higher type 2 diabetes risk (per-allele odds ratio, 1.04; 95% confidence interval, 1.02, 1.05; p=1.1×10^-08^), comparable to previous reports^20^. Considering eleven phenotypes related to glucose homeostasis and metabolic health^21–23^, the A-allele of rs17681684 was significantly associated with insulin disposition index (beta=-0.067, p<0.002)^22^, corrected insulin response (beta=-0.061, p<0.004)^22^, glycated haemoglobin 1c (HbA1c) (beta=0.006, p<0.0003)^21^, and body mass index (beta=0.010, p<5.3×10^-9^), in addition to the previously reported positive association with fasting glucose-dependent insulinotropic peptide (GIP) and the suggestive inverse association with post-glucose load GLP-1 (beta=-0.035, p<4.6×10^-4^)^24^. While sample sizes and hence significance levels for insulin traits were not sufficient to support formal colocalisation analysis, we still obtained a high posterior probability (PP>75%) for a shared genetic signal across plasma citrulline, T2D risk, body mass index, and fasting levels of GIP (**Fig. 5B**). We noted, that the *GLP2R* p.Asp470Asn variant was the only of 6 independent genome-wide significant citrulline-raising loci that was associated with a higher risk of T2D, which indicates that the association does not reflect a general effect of blood citrulline levels on T2D risk but rather a locus-specific association at *GLP2R* (**Fig. 5C**). Plasma citrulline levels have been shown to reflect the volume of intestinal cells and are a marker of GLP2R target engagement in the treatment of short-bowel syndrome with glucagon-like peptide 2 analogues^25^. Taken together, this suggests that genetically higher GLP2R signalling, indicated by the higher citrulline levels among *GLP2R* 470Asn carriers, may lead to chronically elevated GIP (though increased enteroendocrine mass and number of GIP-secreting K-cells), which has been shown to downregulate GIP receptors on pancreatic beta cells^26^, thereby contributing to the observed reduction in the insulin secretory response and increase in T2D risk.

G-protein coupled receptors like GLP2R may signal via G-protein-dependent cyclic adenosine monophosphate (cAMP) production or via G-protein-independent beta-arrestin mediated signalling^27^. To investigate if the *GLP2R* p.Asp470Asn variant affects signalling via either of these pathways, we expressed the *GLP2R* p.Asp470Asn variant in different *in vitro* models (see **Methods**). We show that the variant allele is significantly associated with reduced recruitment of beta-arrestin to GLP2R upon glucagon-like peptide 2 stimulation, but not with cAMP signalling, which suggests a potential role for impaired beta-arrestin recruitment to GLP2R in the pathophysiology of type 2 diabetes (**Fig. 5E-G**).

### Serine and glycine levels play a critical role in the aetiology of a rare eye disease

A recent GWAS of macular telangiectasia type 2 (MacTel), a rare neurovascular degenerative retinal disease, identified three genome-wide susceptibility loci (*PHGDH, CPS1*, and *TMEM161B–LINC00461*) of which the same variants at *PHGDH* and *CPS1* were associated with levels of the amino acids serine and glycine in this GWAS^28^. More recently, it was shown that low serine availability is linked to both MacTel as well as hereditary sensory and autonomic neuropathy type 1 through elevated levels of atypical deoxyshingolipids^29^. Whether genetic predisposition to low serine and glycine levels affects MacTel more generally or has predictive utility has not been investigated. To test this and to explore the specificity of associations between genetic influences on metabolite levels and the risk of MacTel, we generated genetic scores (GS) using the sentinel variants for each of the 141 metabolites with at least one significantly associated locus identified in this GWAS and tested their associations with the risk of MacTel. GS’s for serine and glycine were the only scores associated with risk for MacTel after removal of the known highly pleiotropic *GCKR* variant (**Fig. 6A**). Each standard deviation higher serine levels via the serine GS was associated with a 95% lower risk of MacTel (odds ratio (95% confidence interval), 0.05 (0.03-0.08); p=9.5×10^-30^; **Fig. 6A**). Each of five serine associated variants was individually associated with lower MacTel risk, with a clear dose-response relationship and no evidence of heterogeneity (**Fig. 6B**). The association was unchanged when removing the *GCKR* locus. To disentangle the effect of these two highly correlated metabolites on MacTel risk, we used multivariable Mendelian randomization analysis, which allowed us to test for a causal effect of both measures simultaneously. In this analysis, the effect of serine remained strong, while the effect of glycine was attenuated (**Tab. 2**). Glycine and serine can be interconverted and these results provide genetic evidence that the link between glycine and MacTel is via serine levels through glycine conversion. This hypothesis is supported by the evidence of a log-linear relationship between associations with serine and risk of MacTel among glycine-associated variants (**Fig. 6B**). These findings provide strong evidence that pathways indexed by genetically higher serine levels are strongly and causally associated with protection against MacTel.

**Figure 6.**
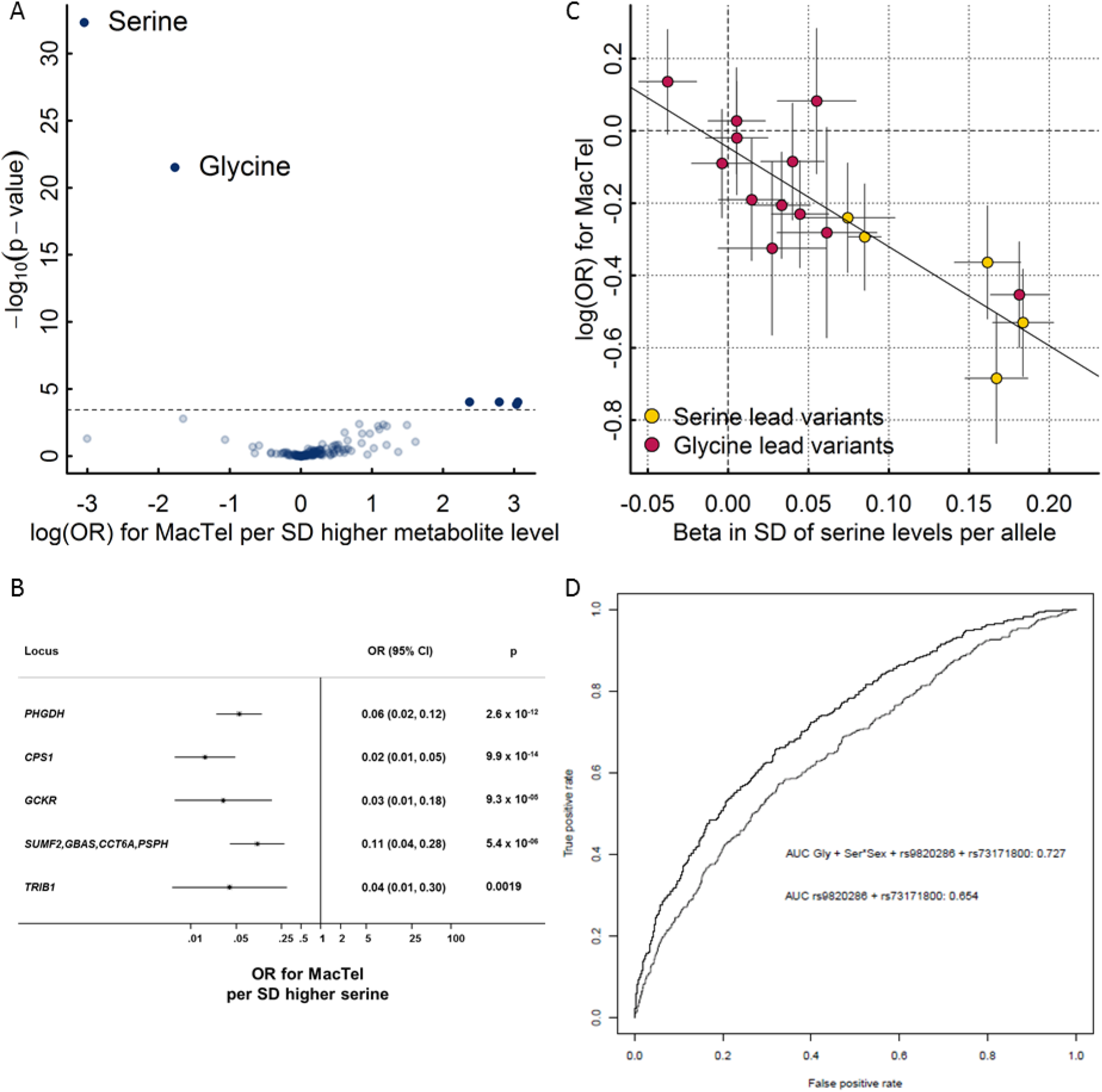
**A** Results from genetic scores for each metabolite on risk for macular telangiectasia type 2 (MacTel). The dotted line indicates the level of significance after correction for multiple testing. The inlet shows the same results but after dropping the pleiotropic variants in *GCKR* and *FADS1-2*. **B** Effect estimates of serine-associated genetic variants on the risk for MacTel. **C** Comparison of effect sizes for lead variants associated with plasma serine levels and the risk for MacTel. **D** Receiver operating characteristic curves (ROC) comparing the discriminative performance for MacTel using a) sex, the first genetic principal component, and two MacTel variants (rs73171800 and rs9820286) not associated with metabolite levels, and b) additionally including genetically predicted serine and glycine at individual levels as described in the methods. The area under the curve (AUC) is given in the legend.

**Table 2.**
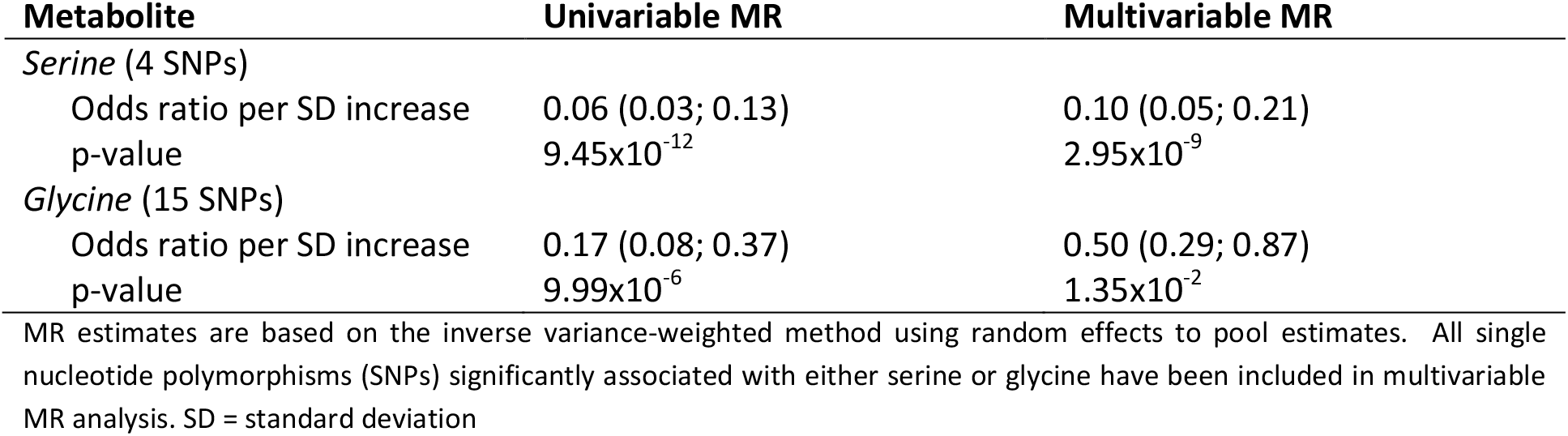
Results from Mendelian randomisation (MR) analysis between metabolite levels and risk of macular telangiectasia type 2.

Given the large observed effect size, we estimated whether using serine and glycine-associated loci might improve the prediction of this rare disease. Adding genetically predicted glycine and serine levels, based on newly discovered metabolite instruments from the present study and previous MacTel variants linked to glycine and serine metabolism, substantially improved prediction of MacTel based on an area under the receiver operating characteristic curve from 0.65 (CI 95%: 0.626-0.682) to 0.73 (0.702-0.753) (**Fig. 6**).

### Common variation at inborn error of metabolism (IEM) associated genes influences the risk of common manifestations of diseases related to the phenotypic presentation of those IEMs

In his seminal 1902 work on alkaptonuria^30^, also known as dark or black urine disease, Archibald Garrod was the first to hypothesise that inborn errors of metabolism are “extreme examples of variations of chemical behaviour which are probably everywhere present in minor degrees”. Previous studies have shown enrichment of metabolite quantitative trait loci in genes known to cause IEMs^31^. Whether or not common variants at IEM causing loci translate into clinically manifest disease remains unknown. The identification of several metabolite-associated variants at IEM-linked genes in this GWAS meta-analysis allows an investigation of the health consequences of genetically determined differences in metabolism for more frequently occurring variants, representing potentially milder forms of the metabolic and other clinical symptoms of IEMs, and providing new candidate genes for rare extreme metabolic disorders that currently lack a genetic basis (**Fig. 7A**). In this study, there were 153 locus-metabolite associations for which 53 unique IEM-associated genes were prioritized as likely causal using either the hypothesis-free genetic approach or the knowledge-based approach on the basis of the Orphanet database^32^. In 89% of these associations (136 of 153) the metabolite associated with a given GWAS locus perfectly matched, or was closely related to, the metabolite affected in patients with the corresponding IEM (**Fig. 7B**).

**Figure 7.**
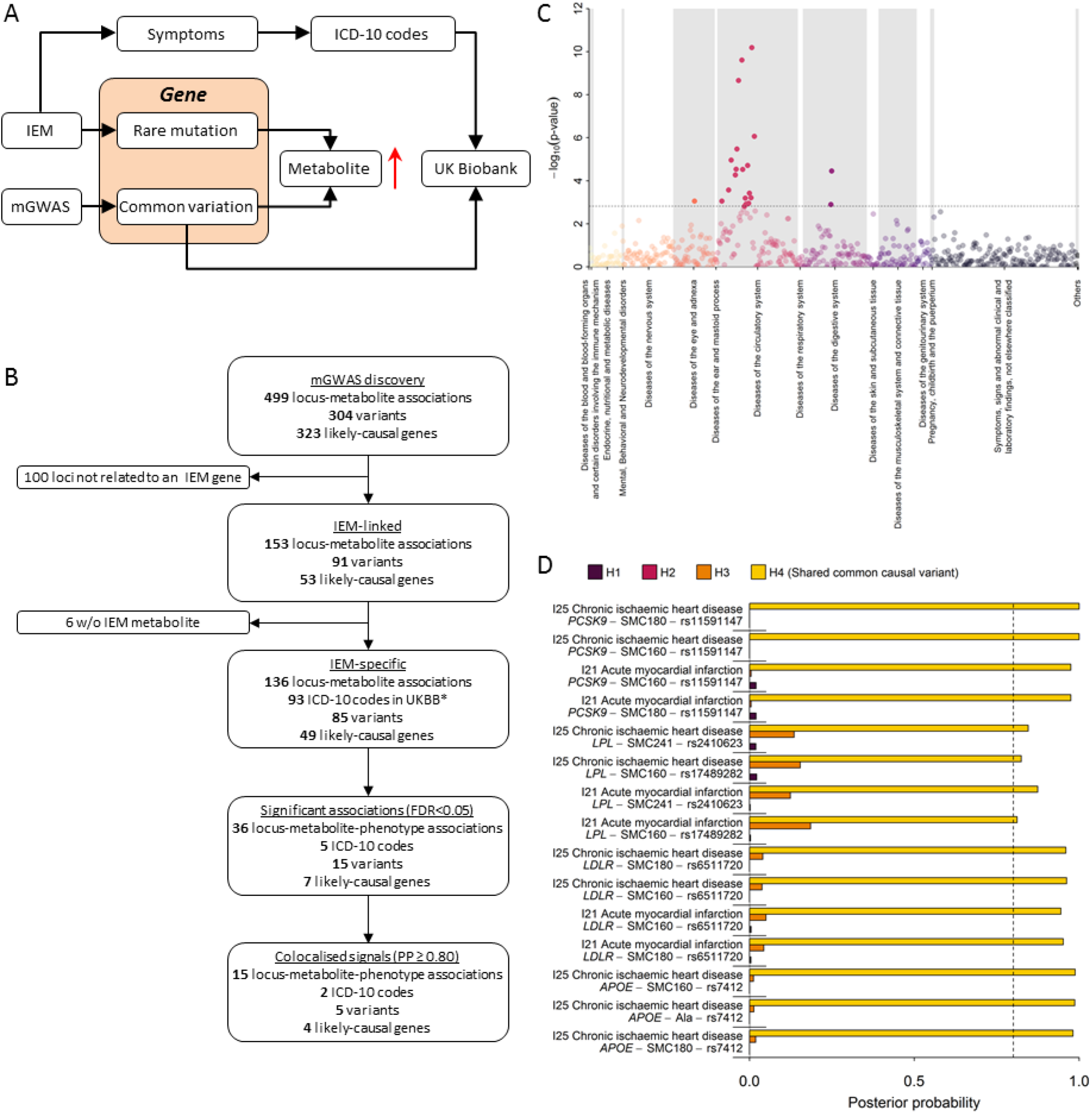
**A** Scheme of the workflow to link common variation in genes causing inborn errors of metabolism (IEM) to complex diseases. **7B** Flowchart for the systematic identification of metabolite-associated variants to genes and diseases related to inborn errors of metabolism (IEM). **C** P-values from phenome-wide association studies among UK Biobank using variants mapping to genes knowing to cause IEMs and binary outcomes classified with the ICD-10 code. Colours indicate disease classes. The dotted line indicates the significance threshold controlling the false discovery rate at 5%. **D** Posterior probabilities (PPs) from statistical colocalisation analysis for each significant triplet consisting of a metabolite, a variant, and a ICD-10 code among UK Biobank. The dotted line indicates high likelihood (>80%) for one of the four hypothesis tested: H0 – no signal; H1 – signal unique to the metabolite; H2 – signal unique to the trait; H3 – two distinct causal variants in the same locus and H4 – presence of a shared causal variant between a metabolite and a given trait.

To test whether IEM-mirroring lead variants from our metabolite GWAS may increase the risk of common manifestations of diseases known to exist in patients with the corresponding IEM (**Fig. 7A**) we obtained a list of electronic health record diagnosis codes (International Statistical Classification of Diseases and Related Health Problems 10th Revision [ICD-10]) and mapped those based on symptoms seen in both, IEM patients and patients with common, complex disease manifestations (see **Methods**). We identified 93 ICD-10 codes with at least 500 cases within the UK Biobank study that aligned with the symptoms or presentations seen in patients with IEMs caused by mutations in genes specifically associated with metabolites observed in the present study. We obtained the association statistics of 85 unique metabolite-associated lead variants at the 136 locus-metabolite associations with these 93 clinical diagnoses and observed 36 associations that met statistical significance (false discovery rate < 5%, **Supplemental Table S6 and Fig. 7B**). For 15 out of those we obtained strong evidence of a shared genetic metabolite-phenotype signal using colocalisation analyses (posterior probability of a shared signal >80%; **Fig. 7D and Supplemental Fig. S3**). These instances linked common genetic variants in or near *APOE, PCSK9, LPL*, and *LDLR* associated with sphingomyelins (SM 16:0, SM 18:0, and SM-OH 24:1) with atherosclerotic heart disease diagnosis codes (I21, I25), mirroring what is observed in rare familial forms of dyslipidaemia in which these sphingomyelins are elevated and the risk of ischemic heart disease is greatly increased^33,34^. These results provide further evidence that common variation at IEM genes can lead to clinical phenotypes and diseases that correspond to those that patients with rare mutations in those same genes are severely affected by. Further studies with detailed follow-up for specific outcomes may provide greater power and help clarify the medical consequences of genetic differences in metabolism caused by metabolite altering variants in the general population.

## Discussion

This large-scale genome-wide meta-analysis has integrated genetic associations for 174 metabolites across different measurement platforms, an approach that has resulted in a three-fold increase in our knowledge of genetic loci regulating levels of these metabolites. We assign likely causal genes for many of the identified associations using a dual approach that combined automated database mining with manual curation.

Previous platform-specific genetic studies of blood metabolites have been substantially smaller in size due to being restricted to a single platform and/ or study^2–10^. We build on these earlier studies to identify and demonstrate enrichment of rare and low-frequency coding variants in enzyme and transporter genes with large effects and reveal the importance of non-linear associations at several loci.

Our results not only provide detailed insight into the genetic determinants of human metabolism but consider their relevance for disease aetiology and prediction. We explore both locus-specific and polygenic score effects and provide tangible examples with clear translational potential. We discovered a strong link between GLP2R, citrulline metabolism and T2D, and demonstrate that the p.Asp470Asn variant underlying the citrulline and T2D associations leads to significantly reduced recruitment of beta-arrestin to GLP2R in various cellular models, providing an explanation for a possible pathological mechanism of a variant previously predicted to be benign^24^.

The finding that a standard deviation increase in serine levels via a genetic score is associated with 95% lower risk of MacTel shows that genetic differences resulting in very specific metabolic consequences can have profound effects on health. Our results suggest that inclusion of genetic scores for metabolite levels can improve identification of high risk individuals. Serine and glycine supplementation and/ or pharmacologic modulation of serine metabolism may help to reduce development or alter the prognosis of this rare, severe eye disease, specifically if targeted to people genetically with a genetic susceptibility to low serine levels. It is important to note, that randomized control trials are needed testing this hypothesis before any recommendations on supplementations could be made.

We finally show specific examples where common genetic variation in IEM-related genes is associated with phenotypes that are also caused by rare highly penetrant mutations. These results suggest that rare variants in metabolite regulating genes newly identified in our study may be valuable candidate genes in patients without a genetic diagnosis but severe alterations in the corresponding or related metabolites. Hence these results provide a new starting point for further investigations into the relationships between human metabolism and common and rare disorders.

## Methods

### Study design and participating cohorts

We performed genome-wide meta-analyses of the levels of 174 metabolites from 7 biochemical categories (amino acids, biogenic amines, acylcarnitines, phosphatidylcholines, lysophosphatidylcholines, sphingomyelins, and sum of hexoses) captured by the Biocrates p180 kit measured using mass spectrometry (MS). As described in more detail below, a total of 174 metabolites were successfully measured in up to 9,363 plasma samples from genotyped participants of the Fenland study^35^.

To maximise sample size and power, we meta-analysed genome-wide association (GWAS) results from the Fenland cohort with those run in the EPIC-Norfolk ^36^ and INTERVAL ^37^ studies, in which metabolites were profiled using MS (Metabolon Discovery HD4 platform) or protein nuclear magnetic resonance (^1^H-NMR) spectrometry ^38 39^ (**Supplementary Tab. 1**). Ten of the 174 Biocrates metabolites were covered across all platforms, while 38 were available on the Biocrates and Metabolon platforms and 126 were unique to Biocrates (**Fig. 1**). We integrated publicly available summary statistics from genome-wide meta-analyses of the same metabolites measured using MS (with Biocrates or Metabolon platforms) or ^1^H-NMR spectrometry (**Supplementary Tab. 1**). Metabolites were matched across platforms by comparing metabolite names and biochemical formulas. Mapping across different Metabolon platforms was done based on retention time/index (RI), mass to charge ratio (m/z), and chromatographic data (including MS/MS spectral data). Scientists at Metabolon Inc. independently reviewed and confirmed metabolite matches.

A summary of the characteristics of participating cohorts is given in **Supplemental Table S1**. The Fenland study is a population-based cohort study of 12,435 participants without diabetes born between 1950 and 1975 ^35^. Participants were recruited from general practice surgeries in Cambridge, Ely and Wisbech (United Kingdom) and underwent detailed metabolic phenotyping and genome-wide genotyping. Ethical approval for the Fenland study was given by the Cambridge Local Ethics committee (ref. 04/Q0108/19) and all participants gave their written consent prior to entering the study. The European Prospective Investigation of Cancer (EPIC)-Norfolk study is a prospective cohort of 25,639 individuals aged between 40 and 79 and living in the county of Norfolk in the United Kingdom at recruitment ^36^. The study was approved by the Norfolk Research Ethics Committee (REC ref. 98CN01) and all participants gave their written consent before entering the study. INTERVAL is a randomised trial of approximately 50,000 whole blood donors enrolled from all 25 static centres of NHS Blood and Transplant, aiming to determine whether donation intervals can be safely and acceptably decreased to optimise blood supply whilst maintaining the health of donors^37^. All participants of the study gave written informed consent and the study was approved by NRES Committee East of England - Cambridge East (ref. 11/EE/0538).

### Metabolomics measurements

The levels of 174 metabolites were measured in the Fenland study by the AbsoluteIDQ^®^ Biocrates p180 Kit (Biocrates Life Sciences AG, Innsbruck, Austria) as reported elsewhere in detail^39,40^. We used a Waters Acquity ultra-performance liquid chromatography (UPLC; Waters ltd, Manchester, UK) system coupled to an ABSciex 5500 Qtrap mass spectrometer (Sciex ltd, Warrington, UK). Samples were derivatised and extracted using a Hamilton STAR liquid handling station (Hamilton Robotics Ltd, Birmingham, UK). Flow injection analysis coupled with tandem mass spectrometry (FIA-MS/MS) using multiple reaction monitoring (MRM) in positive mode ionisation was performed to measure the relative levels of acylcarnitines, phosphatidylcholines, lysophosphatidylcholines and sphingolipids. The level of hexose was measured in negative ionisation mode. Ultra-performance liquid chromatography coupled with tandem mass spectrometry using MRM was performed to measure the concentration of amino acids and biogenic amines. The chromatography consisted of a 5-minute gradient starting at 100% aqueous (0.2% Formic acid) increasing to 95% acetonitrile (0.2% Formic acid) over a Waters Acquity UPLC BEH C18 column (2.1 x 50 mm, 1.7 μm, with guard column). Isotopically labelled internal standards are integrated within the Biocrates p180 Kit for quantification. Data was processed in the Biocrates Met*IDQ* software. Raw metabolite readings underwent extensive quality control procedures. Firstly, we excluded from any further analysis metabolites for which the number of measurements below the limit of quantification (LOQ) exceeded 5% of measured samples. Excluded metabolites were carnosine, dopamine, putrescine, asymmetric dimethyl arginine, dihydroxyphenylalanine, nitrotyrosine, spermine, sphingomyelines SM(22:3), SM(26:0), SM(26:1), SM(24:1-OH), phosphatidylcholine acyl-alky 44:4, and phosphatidylcholine diacyl C30:2. Secondly, in samples with detectable but not quantifiable peaks, we assigned random values between 0 and the run-specific LOQ of a given metabolite. Finally, we corrected for batch-effects with a “location-scale” approach, i.e. with normalization for mean and standard deviation of batches.

The levels of up to 38 metabolites were measured in EPIC-Norfolk and INTERVAL using the Metabolon HD4 Discovery platform. Measurements were carried out using MS/MS instruments. For these measurements, instrument variability, determined by calculating the median relative standard deviation, was of 6%. Data Extraction and Compound Identification: raw data was extracted, peak-identified and quality control-processed using Metabolon’s hardware and software. Compounds were identified by comparison to library entries of purified standards or recurrent unknown entities. Metabolon maintains a library, based upon authenticated standards, that contains the retention time/index (RI), mass to charge ratio (m/z), and chromatographic data (including MS/MS spectral data) of all molecules present in the library. Identifications were based on three criteria: retention index, accurate mass match to the library +/- 10 ppm, and the MS/MS forward and reverse scores between the experimental data and authentic standards. Metabolite Quantification and Data Normalization: Peaks were quantified using area-under-the-curve. A data normalization step was performed to correct variation resulting from instrument inter-day tuning differences. Essentially, each compound was corrected in run-day blocks by registering the medians to equal one (1.00) and normalizing each data point proportionately (termed the “block correction”).

The serum levels of 230 metabolites were measured in the INTERVAL study using ^1^H-NMR spectroscopy^38,41^. Among those, 10 metabolites (creatinine, alanine, glutamine, glycine, histidine, isoleucine, leucine, valine, phenylalanine, and tyrosine) overlapped with what is captured by the Biocrates p180 Kit and were used in the present study. Further details of the ^1^H-NMR spectroscopy, quantification data analysis and identification of the metabolites have been described previously^38,42^. Participants with >30% of metabolite measures missing and duplicated individuals were removed. Metabolite data more than 10 SD from the mean was also removed.

### GWAS and meta-analysis

In Fenland and EPIC-Norfolk, metabolite levels were natural log-transformed, winsorised to five standard deviations and then standardised to a mean of 0 and a standard deviation of 1. Genotypes were measured using Affymetrix Axiom or Affymetrix SNP5.0 genotyping arrays. In brief, genotyping in Fenland was done in two waves including 1,500 (Affymetrix SNP5.0) and 9,369 (Affymetrix Axiom) participants and imputation was done using IMPUTE2 to 1000 Genomes Phase 1v3 (Affymetrix SNP5.0) or phase 3 (Affymetrix Axiom) reference panels (**Supplemental Tab. S1**). Plasma metabolite and genotype data was available for 8,714 (Affymetrix Axiom) and 1,022 (Affymetrix SNP5.0) unrelated individuals. In EPIC-Norfolk, 21,044 samples were forwarded to imputation using 1000 Genomes Phase 3 (Oct. 2014) reference panels (**Supplemental Tab. S1**). Imputed SNPs with imputation quality score less than 0.3 or minor allele account less than 2 were removed from the imputed dataset. Genome-wide association analyses were carried out using BOLT-LMM v2.2 adjusting for age, sex, and study-specific covariates in mixed linear models. Alternatively (when the BOLT-LMM algorithm failed due to heritability estimates close to zero or one) analyses were performed using SNPTEST v2.4.1 in linear regression models, additionally adjusting for the top 4 genetic ancestry principal components and excluding related individuals (defined by proportion identity-by-descent calculated in Plink^43^ > 0.1875 as recommended^44^). GWAS analyses in Fenland were performed within genotyping chip, and associations meta-analysed.

In INTERVAL, genotyping was conducted using the Affymetrix Axiom genotyping array. Standard quality control procedures were conducted prior to imputation. The data were phased and imputed to a joint 1000 Genomes Phase 3 (May 2013)-UK10K reference imputation panel. After QC, a total of 40,905 participant remained with data obtained by ^1^H-NMR spectroscopy. For variants with a MAF of >1% and imputed variants with an info score of >0.4 a univariate GWAS for each of the ten metabolic measures was conducted, after adjustment for technical and seasonal effects, including age, sex, and the first 10 principal components, and rank-based inverse normal transformation. The association analyses were performed using BOLT-LMM v2.2 and R. Data based on the Metabolon HD4 platform was available for 8,455 participants. Prior to the Metabolon HD4 genetic analysis, genetic data were filtered to include only variants with a MAF of >0.01% and imputed variants with an info score of >0.3. Phenotype residuals corrected for age, gender, metabolon batch, INTERVAL centre, plate number, appointment month, the lag time between the blood donation appointment and sample processing, and the first 5 ancestry principal components were calculated for each metabolite and the residuals were standardised prior to the genetic analyses in SNPTEST v2.5.1.

For all GWAS analysis within Fenland, EPIC-Norfolk and INTERVAL, variants with Hardy-Weinberg equilibrium p<1×10^-6^ and associations with absolute value of effect size >5 or standard error (SE) >10 or <0 were excluded; insertions and deletions were excluded.

For each metabolite, we performed a meta-analysis of z-scores (betas divided by standard errors) as a measure of association, signals and loci (see below), using METAL software. Heterogeneity between studies for each association was estimated by Cochran’s Q-test. For each metabolite, we also performed a meta-analysis of beta and standard errors for the subset of studies (Fenland and, when available, EPIC-Norfolk and/or INTERVAL) where we had access to individual level data and standardised phenotype preparation to estimate effect sizes. Quality filters implemented after meta-analysis included exclusion of SNPs not captured by at least 50% of the participating studies and 50% of the maximum sample size for that metabolite and variants with a minor allele frequency below 0.5%. As a result, meta-analyses assessed the associations of up to 13.1 million common or low-frequency autosomal SNPs. Chromosome and base pair positions are determined referring to GRCh37 annotation. To define associations between genetic variants and metabolites, we corrected the conventional threshold of genome wide significance for 102 tests (i.e. p<4.9×10^-10^), corresponding to the number of principal components explaining 95% of the variance of the 174 metabolites in the Fenland cohort, as previously described^45^.

### Signal selection

For each metabolite, we ranked associated SNPs (p <4.9×10^-10^) by z-score to select trait-sentinel SNPs and defined an “association” region as the region extending 1 Mb to each side of the trait-sentinel SNP. During forward selection of trait-sentinel SNPs and loci for each trait, adjacent and partially overlapping association regions were merged by extending region boundaries to a further 1 Mb. After defining trait-sentinel SNPs and association regions we defined overall lead-sentinel SNP and loci for any metabolite using a similar approach. Trait-sentinel SNPs were sorted by z-score for the forward selection of lead-sentinel SNPs and a “locus” was defined as the region extending 1 Mb each side of the lead-sentinel SNP. Regions larger than 2 Mb defined in the trait-sentinel association region definition were carried over in the definition of lead-sentinel SNP loci. As a result, all lead-sentinel SNPs were >1Mb apart from each other and had very low or no linkage disequilibrium (R^2^ < 0.05).

For a given locus, independent signals across metabolites were determined based on linkage disequilibrium (LD)-clumping of SNPs that reached the Bonferroni corrected p-value. SNPs with the smallest p-values and an R^2^ less than 0.05 were identified as independent signals. LD patterns were estimated with SNP genotype data imputed using the haplotype reference consortium (HRC) reference panel, with additional variants from the combined UK10K plus 1000 Genomes Phase 3 reference panel in the EPIC-Norfolk study (n = 19,254 after removing ancestry outliers and related individuals).

Throughout the manuscript, the term “locus” indicates a genomic region (≥1 Mb each side) of a lead-sentinel SNP harbouring one or more trait-sentinel SNPs; “signal” indicates a group of trait-sentinel SNPs in LD with each other but not with other trait-sentinel SNPs in the locus (R^2^ < 0.05); “association” indicates trait-sentinel SNP to metabolite associations defined by a trait-lead SNP and its surrounding region (≥1 Mb each side).

We tested at each locus for conditional independent variants using exact stepwise conditional analysis in the largest Fenland sample (n = 8,714) using SNPTEST v2.5 with the same baseline adjustment as in the discovery approach. To refine signals at those loci we used a more recent imputation for this analysis based on the HRC v1 reference panel and additional SNPs imputed using UK10K and 1000G phase 3. We defined secondary signals as those with a conditional p-value < 5×10^-8^. To avoid problems with collinearity we tested after each round if inclusion of a new variant changed associations of all previous variants with the outcome using a joint model. If this model indicated that one or more of the previously selected variants dropped below the applied significance threshold we stopped the procedure, otherwise we repeated this procedure until no further variant met the significance threshold in conditional models. We considered only locus-metabolite associations meeting the GWAS-threshold for significance in the Fenland analysis (n=228).

### Investigation of heterogeneity

We used a meta-regression model to identify factors associated with larger I^2^ values across all 499 identified SNP-metabolite associations. To this end, a vector of heterogeneity estimates, I^2^, from the meta-analysis was obtained as outcome and the following explanatory variables were considered: strength of effect (absolute Z-score of the SNP – metabolite association), biochemical class, dummy variables indicating the study of origin (related to the measurement platform), and the number of contributing studies as an estimate of sample size. A significant effect of any of those terms in a linear regression model was taken to indicate a source of heterogeneity across SNP-metabolite associations and hence identified systematic factors contributing to any observed cross-platform heterogeneity.

### Statistical fine-mapping

We used statistical fine mapping to determine 99%-credible intervals for all independently associated SNPs using the R package ‘corrcoverage’. Briefly, regional summary statistics (betas and standard errors) were converted to approximate Bayes factors as described in Wakefield *et al*.^46^ to calculate the posterior probability (PP) for each variant driving the association. Credible sets are subsequently defined as the ranked list of variants cumulatively covering 99% of the PP to cover the true causal signal. For loci with evidence of independent secondary signals we used GCTA COJO-cond algorithm to generate conditional association statistics conditioning on all other independent signals in the locus. Since the calculation of approximate Bayes factors requires betas and standard errors we used meta-analysis results across studies for which we had access to individual data (Fenland, EPIC-Norfolk, and INTERVAL). However, out of 546 detected signals 473 reached genome-wide significance (p<5×10^-8^) in this smaller subset and we restricted fine-mapping to those associations.

### Muli-trait colocalisation across metabolites

We used hypothesis prioritisation in multi-trait colocalisation (HyPrColoc)^15^ at each of the identified 144 loci 1) to identify metabolites sharing a common causal variant over and above what could be identified in the meta-analysis to increase statistical power, and 2) to identify loci with evidence of multiple causal variants with distinct associated metabolite clusters. Briefly, HyPrColoc aims to test the global hypothesis that multiple traits share a common genetic signal at a genomic location and further uses a clustering algorithm to partition possible clusters of traits with distinct causal variants within the same genomic region. HyPrColoc provides for each cluster three different types of output: 1) a posterior probability (PP) that all traits in the cluster share a common genetic signal, 2) a regional association probability, i.e. that all the metabolites share an association with one or more variants in the region, and 3) the proportion of the PP explained by the candidate variant. We considered a highly likely alignment of a genetic signal across various traits if the PP > 75% or the regional association probability > 80% and the PP > 50%. The second criterion takes into account that metabolites may share multiple causal variants at the same locus. We used the same set of summary statistics as described for statistical fine-mapping, i.e. based on betas and standard errors across studies for which we had access to individual level data. We further filtered metabolites with no evidence of a likely genetic signal (p>10^-5^) in a region before performing HyPrColoc, which improved clustering across traits by minimizing noise. We used the same workflow to test for the alignment of a genetic signal at the *GLPR2* locus using summary statistics from T2D (see below), a meta-analysis for body mass index across GIANT and UK Biobank, plasma GIP, and plasma citrulline.

### Testing for non-linear effects

We tested each of the 499 identified SNP (j) – metabolite (i) pairs for the deviation from an additive linear model by introducing a dummy variable encoding heterozygous carriers (D), i.e. D = 1 if heterozygous and 0 otherwise, in the following regression model:

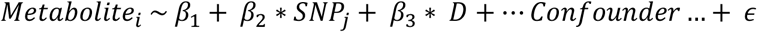

A significant estimate *β*_3_ indicates departure from linearity. In a more formal framework this test allows to test for either a dominant negative or positive model of inheritance depending on the coding of the effect allele. We implemented this test in STATA version 13 using individual level data from the Fenland cohort.

### Metabolic network and community detection

We used Gaussian graphical modelling (GGMs) to construct a metabolic network across all 174 metabolites in a data-driven manner^2^. Briefly, GGMs are based on partial correlation minimizing confounding and have been shown to recover tight biochemical dependencies from single spot blood measurements. The final network comprised 167 metabolites and 554 significant (p<3.3×10^-6^) edges. We next preformed community detection using the Girvan-Newman algorithm, which successively removes edges with high edge betweenness creating a dendrogram of splits of the network into communities, as implemented in the R package *igraph*. We obtained 14 distinct communities including those covering metabolites of distinct biochemical species as well as subdividing larger metabolite classes (**Supplemental Fig. S2**).

### Hypothesis-free (genetic) assignment of causal genes

To assign likely causal genes to lead SNPs at each locus we generated a scoring system. We identified the nearest gene for each variant by querying HaploReg^47^. Next we integrated expression quantitative trait loci (eQTL) studies (GTEx v6p) to identify genes whose expression levels are associated with metabolite levels using TWAS/FUSION (Transcriptome-wide association study / Functional summary-based imputation)^48^. In doing so, we assigned to each variant-metabolite association one or more associated genes using the variant as common anchor. We further assigned higher impact for a causal gene if either the metabolite variant itself or a proxy in high linkage disequilibrium (R^2^>0.8) was a missense variant for a known gene again using the HaploReg database to obtain relevant information. Based on those three criteria we ranked all possible candidate genes and kept those with the highest score as putative causal gene.

### Knowledge-based (biological) assignment of causal genes

Metabolite traits are unique among genetically evaluated phenotypes in that the functional characterization of the relevant genes has often already been carried out using classic biochemical techniques. The objective for the knowledge-based assignment strategy was to find the experimental evidence that has previously linked one of the genes proximal to the GWAS lead variant to the relevant metabolite. For many loci and metabolites this ‘retrospective’ analysis has already been carried out ^31 49^.For these cases, previous causal gene assignments were generally adopted. For novel loci, we employed a dual strategy that combined automated database mining with manual curation. In the automated phase, seven approaches were employed to identify potential causal genes among the 20 protein-coding genes closest to each lead variant, as described in detail below, using the shortest distance determined from the lead SNP to each gene’s transcription start site (TSS) or transcription end site (TES), with a distance value of 0 assigned if the SNP fell between the TSS and TES.

These 7 approaches were as follows:

1. HMDB metabolite names^50^ were compared to each entrez gene name;
2. Metabolite names were compared to the name and synonyms of the protein encoded by each _gene_^51^
3. HMDB metabolite names and their parent terms (class) were compared to the names for the protein encoded by each gene (UniProt).
4. Metabolite names were compared to rare diseases linked to each gene in OMIM^32^ after removing the following non-specific substrings from disease names: uria, emia, deficiency, disease, transient, neonatal, hyper, hypo, defect, syndrome, familial, autosomal, dominant, recessive, benign, infantile, hereditary, congenital, early-onset, idiopathic;
5. HMDB metabolite names and their parent terms were compared to all GO biological processes associated with each gene after removing the following non-specific substrings from the name of the biological process: metabolic process, metabolism, catabolic process, response to, positive regulation of, negative regulation of, regulation of. For this analysis only gene sets containing fewer than 500 gene annotations were retained.
6. KEGG maps^52^ containing the metabolite as defined in HMDB were compared to KEGG maps containing each gene, as defined in KEGG. For this analysis the large “metabolic process” map was omitted.
7. Each proximal gene was compared to the list of known interacting genes as defined in HMDB.

For each text-matching based approach, a fuzzy text similarity metric (pair coefficient) as encoded in the ruby gem “fuzzy_match” was used with a score greater than 0.5 considered as a match.

In the next step, all automated hits at each locus were manually reviewed for plausibility. In addition, other genes at each locus were reviewed if the Entrez gene or UniProt description of the gene suggested it could potentially be related to the metabolite. If existing experimental evidence could be found linking one of the 20 closest genes to the metabolite, that gene was selected as the biologically most likely causal gene. If no clear experimental evidence existed for any of the 20 closest protein coding genes, no causal gene was manually selected. In a few cases multiple genes at a locus had existing experimental evidence. This frequently occurs in the case of paralogs with similar molecule functions. In these cases, all such genes were flagged as likely causal genes.

For each manually selected causal gene, the earliest experimental evidence linking the gene (preferably the human gene) to the metabolite was identified. The median publication year for the identified experimental evidence was 2000.

### Enrichment of type 2 diabetes associations among metabolite associated lead variants

We examined whether the set of independent lead metabolite associated variants (N=168) were enriched for associations with type 2 diabetes. We plotted observed versus expected −log10(p values) for the 168 lead variants in a QQ-plot, using association statistics from a type 2 diabetes meta-analysis including 80,983 cases and 842,909 non-cases from the DIAMANTE study ^53^ (55,005 T2D cases, 400,308 non-cases), UK Biobank^54^ (24,758 T2D cases, 424575 non-cases, application number 44448) and the EPIC-Norfolk study (additional T2D cases not included in DIAMANTE study: 1,220 T2D cases and 18,026 non-cases). This QQ-plot was compared to those for 1000 sets of variants, where variants in each set were matched to the index metabolite variants in terms of MAF, the number of variants in LD (R^2^>0.5), gene density and distance to nearest gene (for all parameters +/- 50% of the index variant value), but otherwise randomly sampled from across the autosome excluding the HLA region. MAF and LD parameters for individual variants were determined from the EPIC-Norfolk study (using the combined HRC, UK10K and 1000G imputation as previously described) and gene information was derived from GENCODE v19 annotation^55^. A one-tailed Wilcoxon rank sum test was used to compare the distribution of association −log10 p-values for the metabolite associated variants with that for the randomly sampled, matched, variants.

### Functional characterisation of D470N mutant GLP2R

To investigate the functional differences between wild-type (WT) GLP2R and the D470N mutant GLP2R we generated D470N GLP2R mutant constructs using site-directed mutagenesis and characterised canonical GLP2R signalling pathways via cAMP as well as alternative signalling pathways via β-arrestin and P-ERK.

#### Generation of D470N GLP2R mutant expressing constructs

Human GLP2R cDNA within the pcDNA3.1+ vector was purchased, and Gibson cloning was completed to insert an internal ribosome entry site (IRES) and venus gene downstream of the GLP2R sequence. Following this, QuikChange Lightning site directed mutagenesis was used to perform a single base change from GAC (encoding aspartic acid) to AAC (encoding asparagine) at amino acid position 470 (**Supplemental Fig. 4A-B**). Successful mutagenesis was confirmed by DNA Sanger sequencing (**Supplemental Fig. 4C**), and the successful products were scaled up for use in functional assays. The WT and mutant GLP2R constructs within the pcDNA3.1+ vector were used to assess signalling by cAMP and P-ERK. To determine β-arrestin recruitment using NanoBiT^®^ technology, an alternative vector was required for lower expression of GLP2R, and fusion of GLP2R to the Large BiT subunit of NanoBiT^®^. For this, GLP2R was cloned into the pBiT1.1_C[TK/LgBiT] vector using restriction cloning and ligation. DNA Sanger sequencing was then used for confirmation of successful cloning.

#### Comparison of WT and D470N GLP2R signalling via cAMP

After generation of WT and D470N GLP2R containing constructs, these were used to assess differences in WT and mutant GLP2R signalling. The initial signalling pathway to be assessed was Gαs signalling via cAMP. CHO K1 cells were transiently transfected with WT or mutant GLP2R constructs, then after 16-24 hours were treated with a dose response of GLP-2. cAMP levels were measured following 30 minutes of GLP-2 treatment, in an end-point lysis HitHunter^®^ cAMP assay. The presence of IRES-Venus within the GLP2R expressing vectors allowed transfection efficiency to be determined for each construct. Transfection efficiency was approximately 60-70%, with no differences between the WT and mutant constructs. Comparison of the GLP-2 dose-response in WT and mutant GLP2R expressing cells revealed no significant differences in signalling, with an almost overlapping dose response curve (**Fig. 5E**).

#### Comparison of β-arrestin recruitment to the WT and D470N GLP2R

Both β-arrestin 1 and β-arrestin 2 recruitment were assessed using a Nano-Glo^®^ live cell assay in transiently transfected HEK293 cells. Briefly, the recruitment of β-arrestin to GLP2R brings the large and small BiT subunit of NanoBiT^®^ together, resulting in increased luciferase activity. The top concentrations from the GLP-2 dose response in the cAMP assay (1–100 nmol/l GLP-2) were chosen for stimulation of the GLP2R and observation of β-arrestin recruitment. Both β-arrestin 1 and β-arrestin 2 were recruited to the WT GLP2R upon GLP-2 stimulation, in a dose-dependent manner (**Supplemental Fig. 5a, c**). The maximal luciferase activity for both β-arrestin 1 and β-arrestin 2 recruitment to the mutant GLP2R was significantly decreased when compared to the WT GLP2R, indicating the extent of β-arrestin recruitment was markedly decreased (**Supplemental Fig. 5b, d**). The example traces indicate that neither β-arrestin 1 or β-arrestin 2 were recruited to the mutant GLP2R upon stimulation with 1 nmol/l GLP-2, however the same concentration of GLP-2 induced β-arrestin recruitment to the WT GLP2R. Overall there was a significant decrease in β-arrestin 1 and β-arrestin 2 recruitment to the D470N GLP2R mutant (**Figure 5F-G**).

### Genetic score and Mendelian randomization analysis for macular telangiectasia type 2

For each metabolite a genetic score (GS) was calculated using all variants meeting genome-wide significance and their beta-estimates as weights obtained from the meta-analysis of studies for which individual level data was available. We used fixed-effect meta-analysis to test for the effect of the GS on MacTel risk using the summary statistics from the most recent GWAS. A conservative Bonferroni-correction for the number of tested GS’s was used to declare significance (p<3.5×10^-4^). Sensitivity analyses were performed where the pleiotropic *GCKR* variant was removed.

To test for causality between circulating levels of glycine and serine for MacTel we performed two types of Mendelian randomization (MR) analysis. In a two-sample univariable MR^56^ we tested for an individual effect of serine (n=4 SNPs) or glycine (n=15 SNPs) on the risk of MacTel using independent non-pleiotropic (i.e. the variant in *GCKR*) genome-wide SNPs as instruments. To this end, we used the inverse variance weighted method to pool SNP ratio estimates using random effects as implemented in the R package *MendelianRandomization*. SNP effects on the risk for MacTel were obtained from^28^. To disentangle the individual effect of those two highly correlated metabolites at the same time we used a multivariable MR model^57^ including all SNPs related to serine or glycine (n=15 SNPs). Beta estimates and standard errors for both metabolites and all SNPs were obtained from the summary statistics and mutually used as exposure variables in multivariable MR. Effect estimates were again pooled using a random effect model as implemented in the R package *MendelianRandomization*. This procedure allowed us to obtain causal estimates for both metabolites while accounting for the effect on each other. Estimates can be interpreted as increase in risk for MacTel per 1 SD increase in metabolite levels while holding the other metabolite constant.

To estimate a potential clinical usefulness of the identified variants we constructed two GRS’s for MacTel using a) sex, the first genetic principal component, and the SNPs rs73171800 and rs9820286 which were identified by the MacTel GWAS study^28^ but not found to be related to either glycine or serine in our study and b) all the previous but additionally including genetically predicted serine and glycine at individual levels, via GS, to the model. An interaction between serine and sex at birth was included to reflect the interaction between SNP rs715 and sex as previously identified ^28^. To assess the predictive ability of both models, receiver operating characteristic curves were computed based on prediction values in 1,733 controls and 476 MacTel cases.

### Identification of genes related to inborn errors of metabolism

Biologically or genetically assigned candidate genes were annotated for IEM association using the Orphanet database^32^. Using a binomial two-tailed test, enrichment of metabolic loci was assessed by comparing the annotated list with the full list of 784 IEM genes in Orphanet against a backdrop of 19,817 protein-coding genes^58^. IEM-annotated loci for which the associated metabolite matched or was closely biochemically related to the IEM corresponding metabolite(s) based on IEMBase^59^ were considered further for analysis.

We hypothesised that IEM-annotated loci with metabolite-specific consequences could also have phenotypic consequences similar to the IEM. To test this, we first obtained terms describing each IEM and translated them into IEM-related ICD-10 codes using the Human Phenotype Ontology and previously-generated mappings^60,61^. We obtained association statistics from the 85 IEM SNPs for phenotypic associations with corresponding ICD-codes among UK Biobank restricting to diseases with at least 500 cases (N=93, **Fig. 7B,** http://www.nealelab.is/uk-biobank). We tested locus-disease pairs meeting statistical significance (controlling the false discovery rate at 5% to account for multiple testing) for a common genetic signal with the corresponding locus-metabolite association using statistical colocalisation. Because of the hypothesis-driven nature of the approach, i.e. prior knowledge of the causal gene and metabolite effect for a given IEM, we adopted an FDR-based strategy to account for multiple testing. We further highlight only those examples with strong evidence for a shared genetic signal (see below).

### Colocalisation analyses

We used statistical colocalisation^62^ to test for a shared genetic signal between a metabolite and a disease of interest. We obtained posterior probabilities (PP) of: H0 – no signal; H1 – signal unique to the metabolite; H2 – signal unique to the trait; H3 – two distinct causal variants in the same locus and H4 – presence of a shared causal variant between a metabolite and a given trait. PPs above 80% were considered highly likely. We used p-values and MAFs obtained from the summary statistics with default priors to perform colocalisation.

## Acknowledgement/Funding

M.P. was supported by a fellowship from the German Research Foundation (DFG PI 1446/2-1). C.O. was founded by an early career fellowship at Homerton College, University of Cambridge. L. B. L. W. acknowledges funding by the Wellcome Trust (WT083442AIA). J.G. was supported by grants from the Medical Research Council (MC_UP_A090_1006, MC_PC_13030, MR/P011705/1 and MR/P01836X/1). Work in the Reimann/Gribble laboratories was supported by the Wellcome Trust (106262/Z/14/Z and 106263/Z/14/Z), UK Medical Research Council (MRC_MC_UU_12012/3) and PhD funding for EKB from MedImmune/AstraZeneca. Praveen Surendran is supported by a Rutherford Fund Fellowship from the Medical Research Council grant MR/S003746/1. A. W. is supported by a BHF-Turing Cardiovascular Data Science Award and by the EC-Innovative Medicines Initiative (BigData@Heart). J.D. is funded by the National Institute for Health Research [Senior Investigator Award] [*]. The EPIC-Norfolk study (https://doi.org/10.22025/2019.10.105.00004) has received funding from the Medical Research Council (MR/N003284/1 and MC-UU_12015/1) and Cancer Research UK (C864/A14136). The genetics work in the EPIC-Norfolk study was funded by the Medical Research Council (MC_PC_13048). Metabolite measurements in the EPIC-Norfolk study were supported by the MRC Cambridge Initiative in Metabolic Science (MR/L00002/1) and the Innovative Medicines Initiative Joint Undertaking under EMIF grant agreement no. 115372. We are grateful to all the participants who have been part of the project and to the many members of the study teams at the University of Cambridge who have enabled this research. The Fenland Study is supported by the UK Medical Research Council (MC_UU_12015/1 and MC_PC_13046). Participants in the INTERVAL randomised controlled trial were recruited with the active collaboration of NHS Blood and Transplant England (www.nhsbt.nhs.uk), which has supported field work and other elements of the trial. DNA extraction and genotyping was co-funded by the National Institute for Health Research (NIHR), the NIHR BioResource (http://bioresource.nihr.ac.uk) and the NIHR [Cambridge Biomedical Research Centre at the Cambridge University Hospitals NHS Foundation Trust] [*]. Nightingale Health NMR assays were funded by the European Commission Framework Programme 7 (HEALTH-F2-2012-279233). Metabolon Metabolomics assays were funded by the NIHR BioResource and the National Institute for Health Research [Cambridge Biomedical Research Centre at the Cambridge University Hospitals NHS Foundation Trust] [*]. The academic coordinating centre for INTERVAL was supported by core funding from: NIHR Blood and Transplant Research Unit in Donor Health and Genomics (NIHR BTRU-2014-10024), UK Medical Research Council (MR/L003120/1), British Heart Foundation (SP/09/002; RG/13/13/30194; RG/18/13/33946) and the NIHR [Cambridge Biomedical Research Centre at the Cambridge University Hospitals NHS Foundation Trust] [*].The academic coordinating centre would like to thank blood donor centre staff and blood donors for participating in the INTERVAL trial.

This work was supported by Health Data Research UK, which is funded by the UK Medical Research Council, Engineering and Physical Sciences Research Council, Economic and Social Research Council, Department of Health and Social Care (England), Chief Scientist Office of the Scottish Government Health and Social Care Directorates, Health and Social Care Research and Development Division (Welsh Government), Public Health Agency (Northern Ireland), British Heart Foundation and Wellcome.

*The views expressed are those of the author(s) and not necessarily those of the NHS, the NIHR or the Department of Health and Social Care.

UK Biobank: This research has been conducted using the UK Biobank resource under

Application Number 44448.

## Author Contribution

Concept and design: L.A.L. and C.L.

Generation, acquisition, analysis and/or interpretation of data: all authors.

Drafting of the manuscript: L.A.L., M.P., and C.L.

Critical review of the manuscript for important intellectual content and approval of the final version of the manuscript: all authors.

## Data Availability

All genome-wide summary statistics will be made available through an interactive webserver upon publication of the manuscript.

## Code Availability

Each use of software programs has been clearly indicated and information on the options that were used is provided in the Methods section. Source code to call programs is available upon request.

