## Supplemental Material for "Cross-platform genetic discovery of small molecule products of metabolism and application to clinical outcomes"

### FIGURES

#### Figure S1

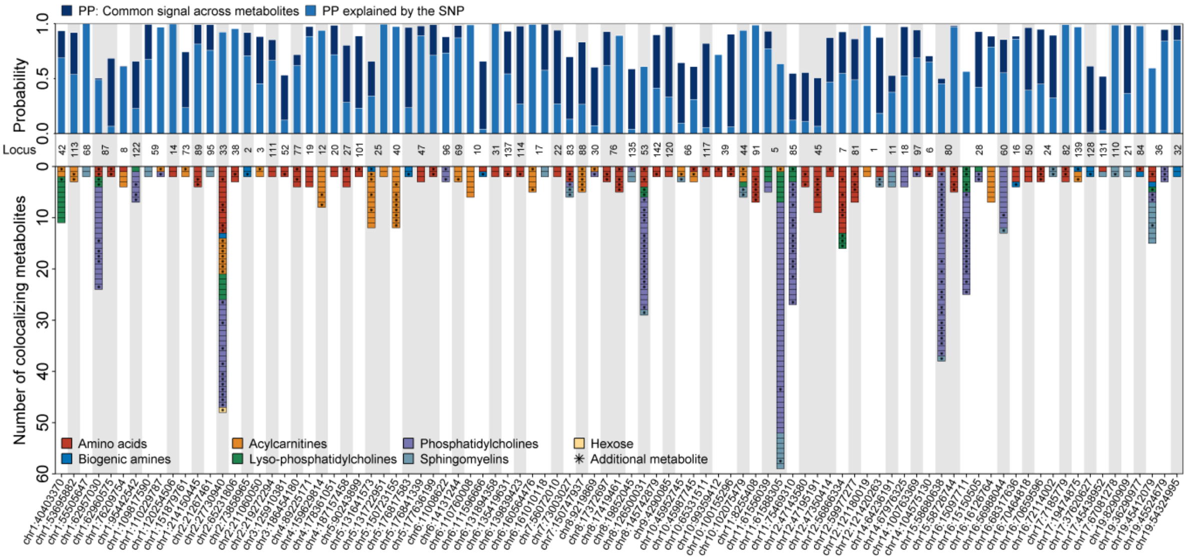


**Figure S1** Summary of statistical colocalization analysis across metabolites for 72 loci with evidence (posterior probability >75% or regional probability >80%) for a shared genetic signal across at least two metabolites. The upper panels display the posterior probabilities (PP) for a shared genetic signal across metabolites (dark blue) and the PP explained by the candidate SNP displayed on the x-axis (blue). The lower panel displays the number and class of metabolites sharing a genetic signal. Metabolites discovered using colocalization only, are indicated by stars. The row in the middle refers to the numbering of loci from Supplemental Table S2.

**Figure S2**

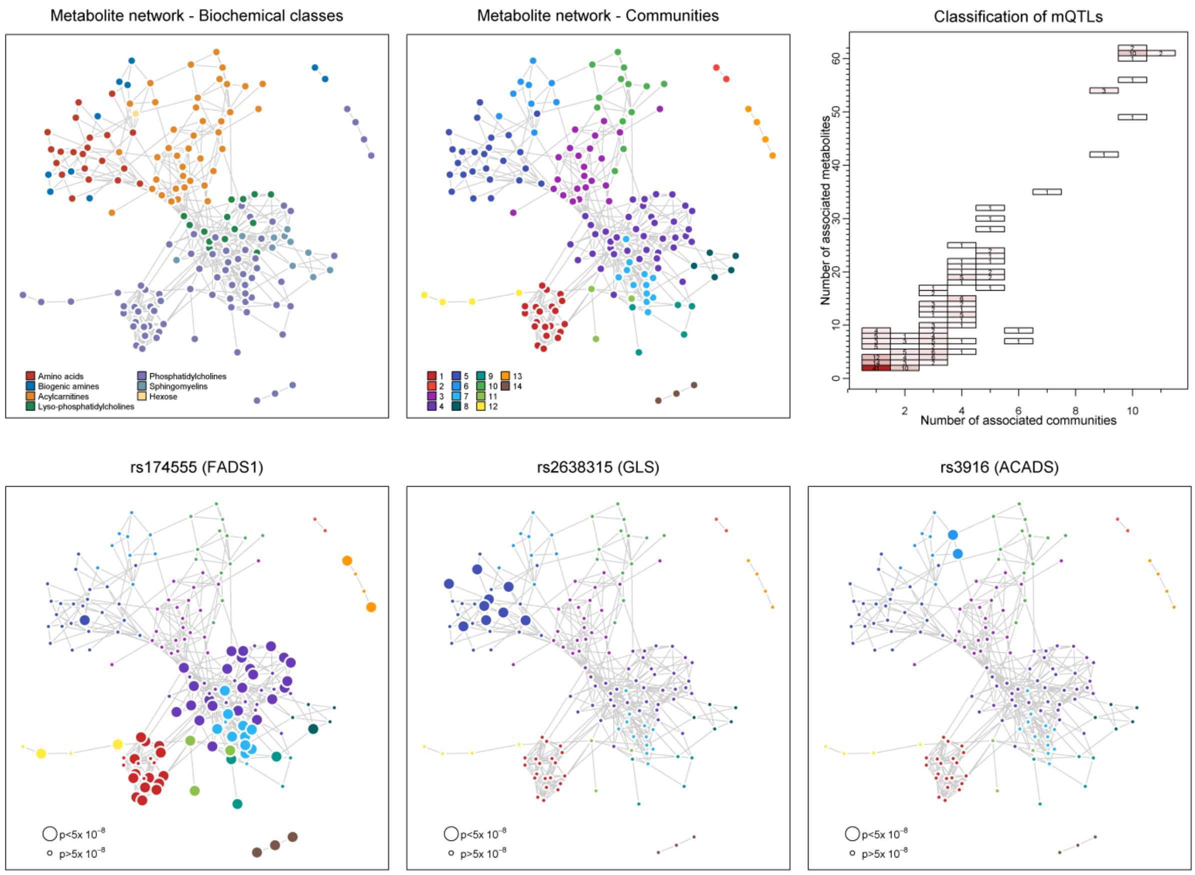


**Figure S2** Derived metabolic network from Gaussian graphical modelling. Each edge represents a significant partial correlation between metabolites, i.e. a significant correlation between two metabolites after taking into account all other metabolites in the network. The left and middle plot in the upper panel show the metabolite network coloured by biochemical class versus identified modules/communities, respectively. The right plot in the upper panel shows the number of associated metabolites ($p<5x10^{-8}$) for each of the 304 metabolite quantitative trait loci against the distribution of those across the number of identified communities. The lower panel maps specific variants onto the network indicating significance by node size.

**Figure S3**

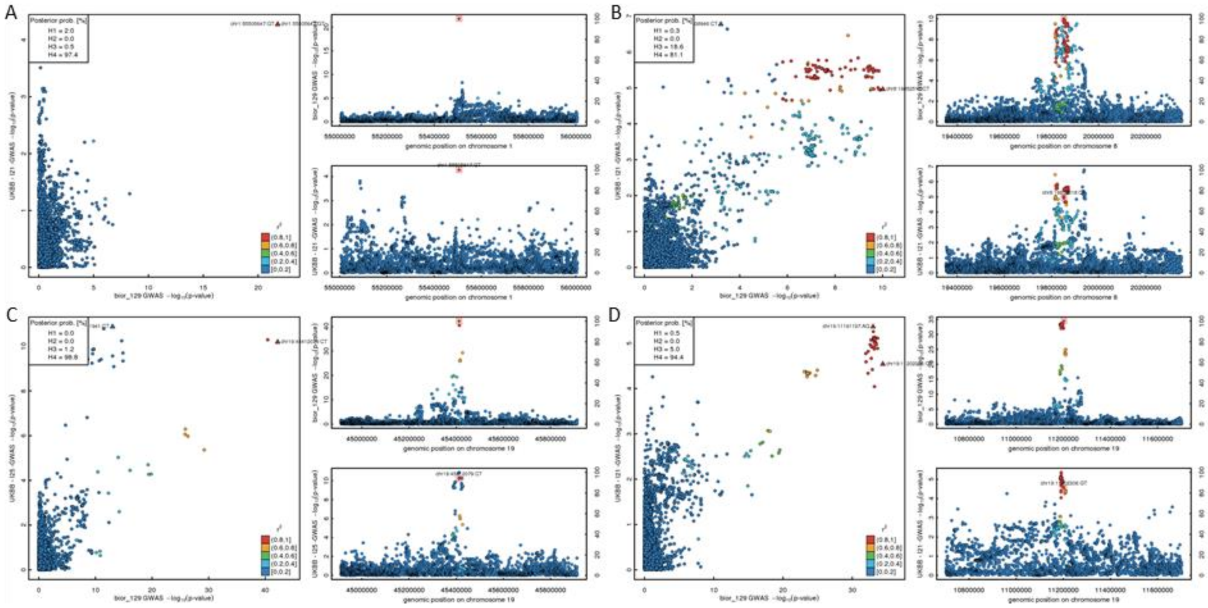


**Figure S3.** Triplet of plots to compare regional associations profiles for results (p-values on a –log10 scale) from genome-wide association analysis (GWAS) for SM C16:0 (x-axis, bior_129) and a disease trait from UK Biobank (y-axis, I21 – Acute myocardial infarction, I25 – Chronic ischaemic heart disease) at A) *PCSK9,* B) *LPL*, C) *LDLR*, and D) *APOE*. Regions were chosen based on the purification workflow to reveal links to inborn errors of metabolism (see Main text). The legend displays posterior probabilities from statistical colocalisation analyses on the following hypothesis: H0 – no signal; H1 – signal unique to the metabolite; H2 – signal unique to the trait; H3 – two distinct causal variants in the same locus and H4 – presence of a shared causal variant between a metabolite and a given trait. The strongest variants for each trait in the locus are annotated and LD-based colouring was done with respect to the lead variant.

**Figure S4**

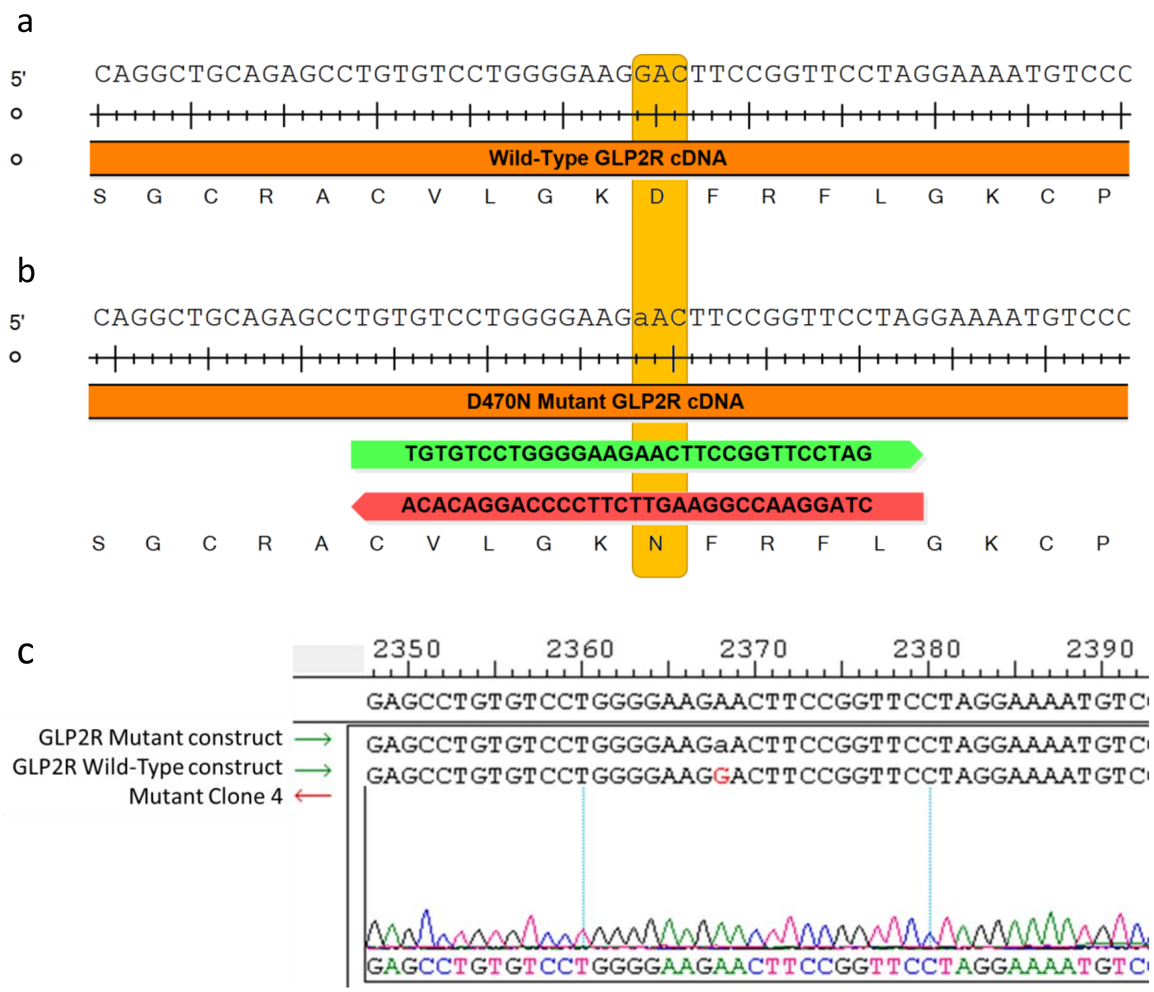


**Figure S4. Schematic depicted wild-type and mutant GLP2R sequences.** (a) Wild-type GLP2R sequence, amino acid targeted for mutation is highlighted in yellow. (b) Mutant GLP2R sequence, showing D470N mutation highlighted in yellow. Primers used to introduce the mutation using QuikChange Lightning Site-Directed Mutagenesis are depicted, forward primer shown in green, and reverse primer shown in red. (c) Sequence confirmation of successfully mutatgenesis, chromatogram depicted for clone 4 which was then sequenced in full.

**Figure S5**

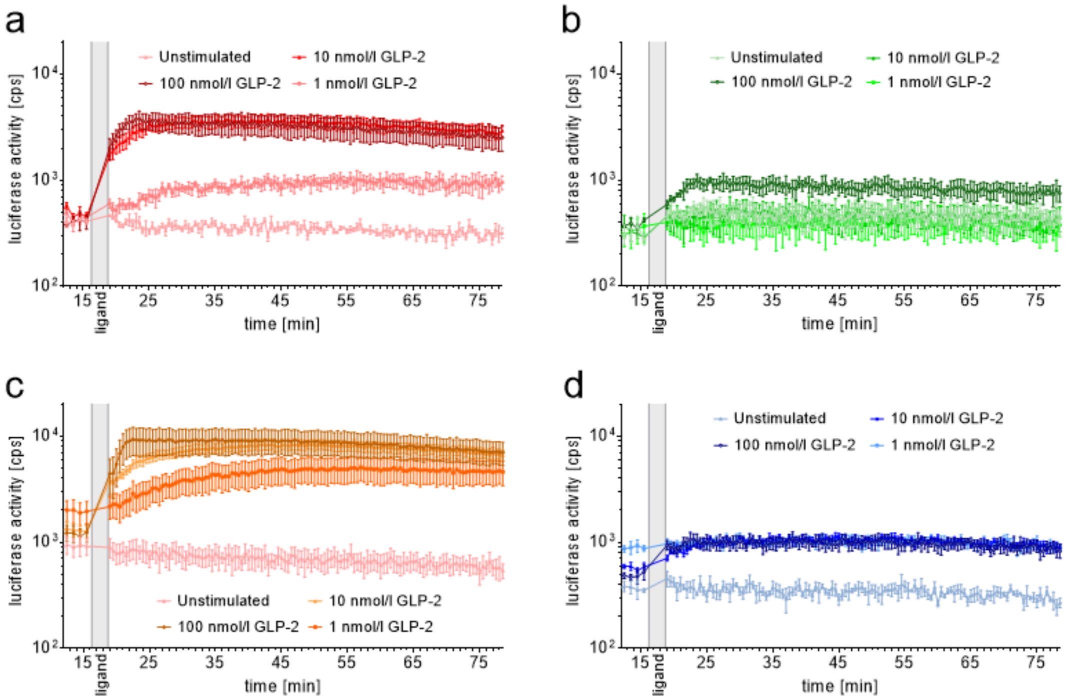


**Figure S5. Example traces of GLP2R wild-type and mutant signalling via beta-arrestin 1 and beta-arrestin 2.** Example real-time beta arrestin responses to a dose titration of GLP-2 are displayed for beta-arrestin 1 (a, b) for wild-type GLP2R (a) and D470N mutant GLP2R (b). Real time beta arrestin responses are also displayed for beta-arrestin 2 (c, d) for wild-type GLP2R (c) and D470N mutant GLP2R (d). Data are from representative experiments, and are displayed as mean ± SEM from 3 wells.
